# Spatial and feature-selective attention interact to drive selective coding in frontoparietal cortex

**DOI:** 10.1101/2024.12.03.626620

**Authors:** Nadene Dermody, Romy Lorenz, Erin Goddard, Arno Villringer, Alexandra Woolgar

**Affiliations:** MRC Cognition and Brain Sciences Unit, University of Cambridge, UK; Max Planck Institute for Biological Cybernetics, Tübingen, Germany; Max Planck Institute for Human Cognitive and Brain Sciences, Leipzig, Germany; University of New South Wales, Sydney, Australia; Berlin School of Mind and Brain, Berlin, Germany; Clinic of Cognitive Neurology, Leipzig University, Leipzig, Germany; Charité University Medicine Berlin, Berlin, Germany; Department of Psychology, University of Cambridge, UK

## Abstract

Attention enables the selective processing of relevant information. Two types of selective attention, spatial and feature attention, have separable neural effects but in real life are often used together. Here, we asked how these types of attention interact to affect information coding in a frontoparietal ‘multiple-demand’ (MD) network, essential for attentional control. Using functional magnetic resonance imaging (fMRI) with multivariate pattern analysis, we examined how covert attention to object features (colour or shape) and spatial locations (left or right) influences coding of task-related stimulus information. We found that spatial and feature attention interacted multiplicatively on information coding in MD and visual regions, such that there was above-chance decoding of the attended feature of the attended object and no detectable coding of visually equivalent but behaviourally irrelevant aspects of the visual display. The attended information had a multidimensional neural representation, with stimulus information (e.g., colour) and discrimination difficulty (distance from the categorical decision boundary) reflected in separate dimensions. Rather than boosting processing of whole objects or relevant features across space, our results suggest neural activity reflects precise tuning to relevant information, indicating a highly selective control process that codes behaviourally relevant information across multiple dimensions.

## 1. Introduction

Everyday experience tells us that there are limits to the amount of information we can process at any one time. To find a friend in a crowded bar, we can ignore aspects of the environment irrelevant to our goal and zero in on what is relevant, focussing on likely locations and paying attention to particular features (e.g., hair colour). Selective attention describes the process by which a subset of task-relevant inputs is ‘selected’ for prioritised processing from the wealth of information available to us. Behaviourally, attention enhances perception of the attentional target relative to unattended stimuli (e.g., see Carrasco, 2011). At the neural level, attention enhances the signal of the attended information (e.g., Duncan, 2001; Noudoost et al., 2010; Reynolds & Heeger, 2009; Thiele & Bellgrove, 2018). For example, in non-human primates, attending to a particular stimulus can increase the response (e.g., firing rate) of neurons responding to that stimulus (e.g., Burrows & Moore, 2009; Luck et al., 1997; Luo & Maunsell, 2018; McAdams & Maunsell, 1999; Thiele et al., 2016), reduce response variability of individual neurons and shared variability across neurons (Cohen & Maunsell, 2009; Kanashiro et al., 2017; Mitchell et al., 2007), and is associated with a reduction in the response of cells tuned to an unattended stimulus (Moran & Desimone, 1985; see Thiele & Bellgrove, 2018 for a review). In humans, attention to a particular spatial location, object or feature is reflected in increased activation in related cortical (e.g., Heinze et al., 1994; O’Craven et al., 1999; Sàenz et al., 2002) and subcortical areas (e.g., O’Connor et al., 2002), and reduced neural variability (Arazi et al., 2019). In addition, multivariate decoding of human neuroimaging data repeatedly shows stronger representation of attended relative to unattended information (e.g., Jackson et al., 2017; Jackson & Woolgar, 2018; Jehee et al., 2011; Keller et al., 2022; Li et al., 2018; Woolgar, Williams, et al., 2015) that is sustained over time (Barnes et al., 2022; Battistoni et al., 2020; Goddard et al., 2022; Grootswagers et al., 2021; Moerel et al., 2022).

The process of prioritising relevant information is thought to depend on frontoparietal cortex (e.g., Miller & Cohen, 2001). In particular, in humans, a network of frontoparietal ‘multiple-demand’ (MD) regions (Duncan, 2010; Duncan & Owen, 2000) is heavily implicated in driving preferential focus on task-relevant information. This collection of strongly interconnected frontoparietal regions is active during a wide range of cognitively demanding tasks (Assem, Blank, et al., 2020; Fedorenko et al., 2013) and has also been referred to as the task positive network (Fox et al., 2005), frontoparietal control network (Vincent et al., 2008), task-activation ensemble (Seeley et al., 2007), or cognitive control network/networks (Cole & Schneider, 2007; Gratton et al., 2018), and is thought to comprise ‘flexible hubs’ that rapidly update their brain-wide functional connectivity patterns in accordance with task demands (Cole et al., 2013). MD regions are thought to adaptively prioritise coding of task-relevant information (Duncan, 2001), biasing processing elsewhere in the brain to drive widespread congruence in representations (Desimone & Duncan, 1995; Miller & Cohen, 2001).

In line with this, MD regions demonstrate flexible coding properties, including the capacity to represent a wide range of different types of task information (Schultz et al., 2022; Woolgar et al., 2016; Zheng et al., 2024) and an adaptive response to changing task demands (e.g., Woolgar, Afshar, et al., 2015; Woolgar et al., 2011; Woolgar, Williams, et al., 2015). Moreover, MD regions have been shown to preferentially represent attended or task-relevant information, with previous work in the visual domain, for instance, demonstrating preferential representation of the attended object or object feature (e.g., Jackson et al., 2017; Jackson & Woolgar, 2018; Woolgar, Williams, et al., 2015). Moreover, brain stimulation studies demonstrate that alpha-band oscillatory activity in right parietal cortex appears to have a causal role in supporting neural representation of where attention should be directed in space (Lu et al., 2024), while activity in right dorsolateral prefrontal cortex has been shown to have a causal role in supporting coding of task-relevant object features (Jackson et al., 2021).

In these studies, attention is typically directed to a particular spatial location, target object or target feature, reflecting some of the ways in which visual selective attention can be deployed. These subtypes of selective attention are thought to differentially affect the scope of information prioritised for further processing. For example, spatial attention, whereby attention is directed to a particular location, is associated with selection of all information at the attended location for further processing (e.g., Kastner et al., 1999). Object-based attention theories posit that attention directed to a particular object leads to enhancement of all information within the bounds of an attended object (Duncan, 1984; O’Craven et al., 1999), including multiple features of that object (Duncan, 1984; Egly et al., 1994). In contrast, feature-based attention, whereby attention is directed to a particular feature (e.g., the colour red), is associated with enhanced processing of that feature across the visual field (see Liu, 2019; Maunsell & Treue, 2006 for reviews of related literature), with associated improvements in behavioural performance (e.g., greater detection, discrimination of the attended feature) and more interference from distractors possessing a similar feature, regardless of location (e.g., Rossi & Paradiso, 1995; Sàenz et al., 2003; Zhang & Luck, 2009). Attention can alternatively be directed to an entire feature dimension (e.g., to make fine discriminations of colour while ignoring shape), sometimes known as feature-selective attention, resulting in enhanced separation of task-relevant categories along the attended dimension (X. Chen et al., 2012; Jackson et al., 2017; see also Wisniewski et al., 2023).

Yet in real life, we rarely deploy attention so singularly. Instead, the target of our attention is usually informed by a combination of spatial and feature cues against a backdrop of many other possible competing inputs. Searching for a particular brand of rolled oats in our local supermarket, for example, requires attention direction to the general area in which the oats are typically located (e.g., the bottom shelf of the cereal aisle) and to the most prominent feature of the preferred item (e.g., colour: black, or shape: rectangular box), while ignoring information irrelevant to this task. Despite a wealth of behavioural data on the interaction between different attentional subtypes, there are few neuroimaging studies examining the specificity of task-relevant representations when multiple subtypes of attention are combined.

Guided by seminal theories, an additive effect might be predicted when both spatial and feature attention are deployed, resulting in enhancement of both irrelevant features of attended objects (due to object-based attentional spread; e.g., Brawn & Snowden, 2000; Duncan, 1984; Egly et al., 1994) and relevant features of unattended objects (due to field-wide feature attention effects; e.g., Liu et al., 2007; Sàenz et al., 2002; Treue & Martínez-Trujillo, 1999; Zhang & Luck, 2009), relative to irrelevant features of unattended objects. However, we recently reported a different effect in neural coding, in which there was a striking multiplicative interaction of spatial and feature-selective attention (Goddard et al., 2022). Using multivariate decoding of magnetoencephalography (MEG) signals to track the dynamics of information coding in the human brain, we found that frontal cortex maintained a selective representation of only task-relevant information that was selected by both types of attention (i.e., the task-relevant feature at the selected location). In contrast, information across the visual field was initially decodable in occipital cortex irrespective of attentional condition. Interestingly, the highly tuned response in frontal regions appeared to Granger-cause a sustained selective response in occipital cortex. That is, in occipital cortex, a sustained multiplicative interaction between spatial and feature attention appeared to emerge based on feedback from frontal regions.

In this study, we sought to clarify the spatial basis for this effect using functional magnetic resonance imaging (fMRI). Given the proposed role of the MD system in attentional control, we asked whether the same selective focus on information at the intersection of spatial and feature attention would be seen in the MD system, or whether this system would demonstrate a pattern of information coding more in line with seminal theories of object-based and feature-based attention. Participants completed a covert attention task that manipulated both spatial and feature attention, and we used multivariate pattern analysis (MVPA) to examine the visual stimulus information held in MD regions and elsewhere in the brain, under each attentional condition. We asked whether spatial and feature attention would interact to modulate stimulus information coding in MD regions. In addition, as our paradigm comprised stimuli which varied incrementally across feature space, we sought to characterise how attended information was organised in these regions. We then compared this result to visual cortex, and across the brain using a roaming searchlight.

## 2. Materials and Methods

### 2.1. Participants

Thirty participants (16 female, 14 male, mean age=27.20, SD=4.16) took part in the fMRI experiment. All participants were right-handed native German speakers, had normal or corrected to normal vision, including self-reported normal colour vision. The study was approved by the local ethics committee at the Medical Faculty of the University of Leipzig. All participants gave written informed consent prior to the study and were reimbursed for their time.

### 2.2. Task Design

#### 2.2.1. Stimuli

We used the visual attention paradigm developed by Goddard et al. (2022) with task instructions translated into German. Stimuli comprised a set of 16 novel ‘spiky’ objects which varied along two dimensions (colour and shape, see Figure 1, Panel A). The bounding box for individual objects was 621 pixels (10 degrees of visual angle (dva)) wide by 621 pixels (10 dva) high. Their total size varied with their spikes, but the spikes never reached the border of the object image.

**Figure 1.**
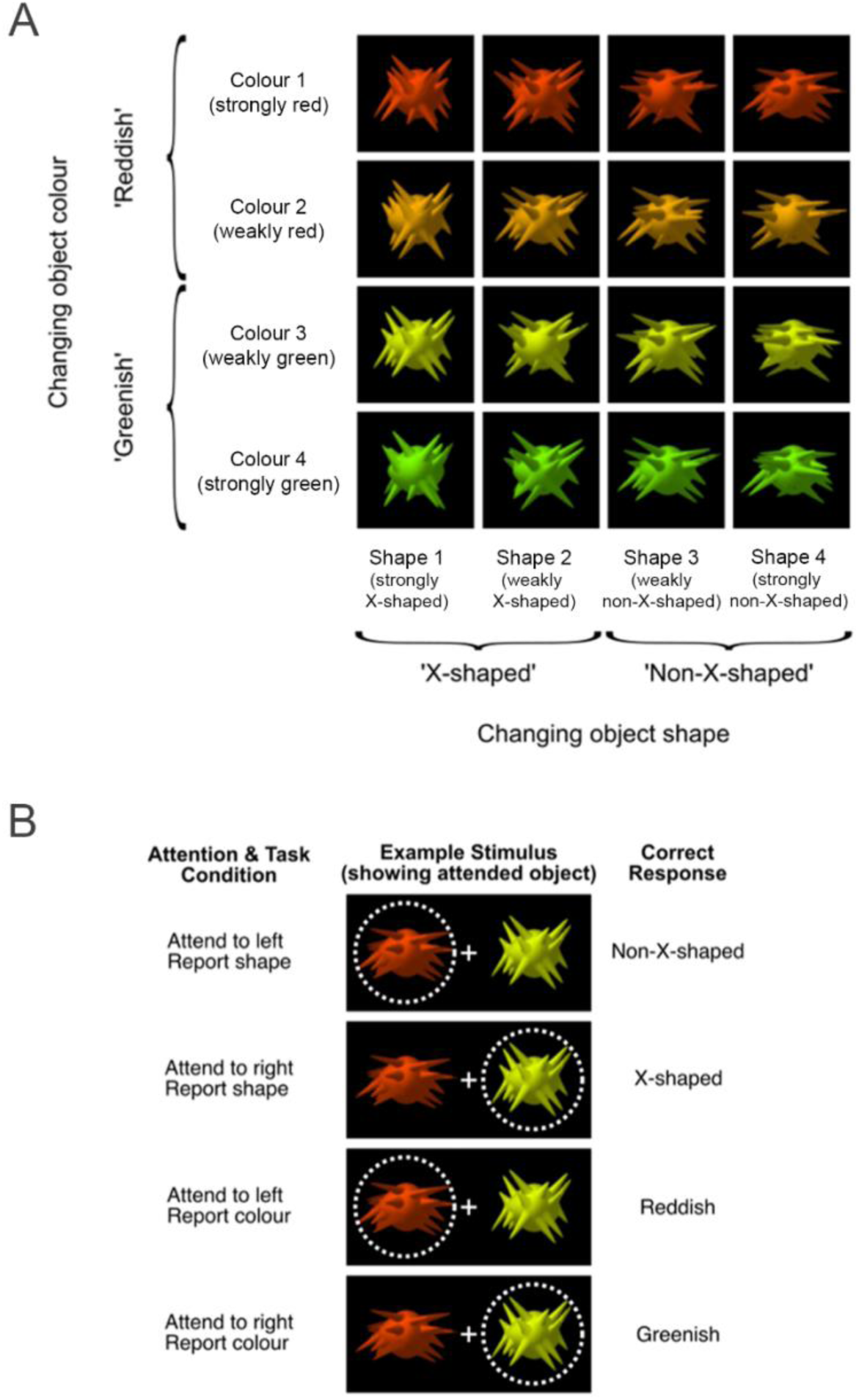
Panel A: Visual stimuli. Each object varied along two dimensions: colour and shape. In the colour task, participants categorised the attended object as either ‘reddish’ or ‘greenish’. In the shape task, participants categorised the attended object as either ‘X-shaped’ or ‘non-X-shaped’, based on the orientations of the object’s spikes. Panel B: Visual stimuli as presented in the scanner. Participants were asked to covertly attend to one side of space, and report (via button press) either the shape or colour of the object in the attended space. The dotted circle shows the locus of spatial attention and was not shown to participants. Figure adapted from Goddard et al. (2022), reproduced under CC BY licence.

The spike orientation statistics were varied to create four shape classes: ‘strongly X-shaped’ (shape 1), ‘weakly X-shaped’ (shape 2), ‘weakly non-X-shaped’ (shape 3), ‘strongly non-X-shaped’ (shape 4). To discourage participants using a single spike to judge shape, each class comprised 100 exemplars with slight variations in location, length, and orientation of the spikes. For the shape-based task, participants classified the target object as ‘X-shaped’ (shape 1 or 2) or ‘non-X-shaped’ (shape 3 or 4, Figure 1, Panel A).

We also varied colour to create four classes: ‘strongly red’ (colour 1), ‘weakly red’ (colour 2), ‘weakly green’ (colour 3), ‘strongly green’ (colour 4). For the colour-based task, participants classified the target object as either ‘reddish’ (colour 1 or 2) or ‘greenish’ (colour 3 or 4, Figure 1, Panel A). The four feature values along each dimension meant that for both tasks, stimuli were either far from the decision boundary (e.g., strongly red: ‘easy’ trials), or close to the decision boundary (e.g., weakly red: ‘hard’ trials). Our intention was that the four colours would be equally spaced along the equiluminant plane joining the strongly red and strongly green coordinates in CIE L’u’v’ space. However, due to technical difficulties with the screen calibration on the projector system, the colours were not fully calibrated for some participants. For the first set of participants (*N*=17), the u’v’ chromaticity coordinates for the colours were as follows: strongly red: 110.34, 41.38, weakly red: 62.07, 48.28, weakly green: 33.10, 52.41, strongly green: -15.17, 58.31. For the remaining participants (*N*=13), luminance was set to 48cd m^2^ for all four colours, with u’v’ chromaticity coordinates as follows: strongly red: 33.03, 28.48, weakly red: 18.21, 30.90, weakly green: 9.31, 32.34, strongly green: -5.52, 34.76. The slight changes in colour between these two sub-groups of participants was included as a predictor in our analyses but did not have a meaningful impact.

#### 2.2.2. Task

On each trial, two stimuli were presented simultaneously against a black screen, one left and one right of a grey fixation cross, with height and width of 1dva (Figure 1, Panel B). Participants were instructed at the beginning of every run to covertly attend to one side of space and report either the colour (‘reddish’ or ‘greenish’) or the shape (‘X-shaped’ or ‘non-X-shaped’) of the attended object. Participants responded by pressing one of two response keys on a button box using the index and middle fingers of their right hand.

#### 2.2.3. Training

To ensure familiarity with the stimulus categories (i.e., ‘reddish’ and ‘greenish’ for colour, and ‘X-shaped’ and ‘non-X-shaped’ for shape) and task instructions, participants completed a training session on a computer located outside the fMRI scanner. Each participant completed a minimum of two training runs, one for the colour task and one for the shape task, each comprising 30 trials. If participants scored below 70% for either the colour or shape task run, they completed additional runs of that task type until their performance reached 70% or above. Training took approximately 20 minutes.

### 2.3. fMRI Acquisition

Functional MRI data were acquired using a Siemens 3.0-Tesla Skyra scanner (Siemens, Erlangen) with 32-channel head coil at the Max Planck Institute for Human Cognitive and Brain Sciences (MPI CBS) in Leipzig, Germany. We used a high resolution interleaved T2*-weighted echo planar imaging (EPI) acquisition sequence covering the whole brain. The following parameters were used: repetition time (TR) 1000ms; echo time (TE) 30ms; multi-band acceleration factor of 3; 45 interleaved slices of 3.0mm slice thickness with a 10% interslice gap; flip angle of 61 degrees; field of view (FOV) 204mm. AutoAlign was used to correct for movement between acquisition runs. We also acquired T1-weighted MPRAGE structural images for all participants (resolution 1.0 x 1.0 x 1.0mm).

### 2.4. Procedure

Stimuli were presented using MATLAB with Psychophysics Toolbox-3 (Brainard, 1997; Kleiner, Brainard, & Pelli, 2007) and were back-projected onto a screen viewed through a head-coil mounted mirror. Participants, lying supine, viewed the screen from a distance of 95cm. Covert attention was ensured using an EyeLink 1000 Plus remote tracking system; participants were required to fixate on the fixation cross for a variable duration before the task would proceed (details as follows). The eye tracker tracked the participant’s right eye.

Participants performed eight runs in total, where each run comprised a unique combination of the four possible spatial and feature attention manipulations (i.e., attend to one of the two locations: left or right, and one of the two features: colour or shape) and the two response mappings (e.g., button one for ‘reddish’, button two for ‘greenish’). Run order was counterbalanced across participants and arranged such that the attended location and attended feature alternated between runs. Starting location and attended feature were also counterbalanced across participants. Each participant’s imaging data were collected in a single session of approximately 120 minutes.

At the start of each run, participants first completed an eye tracker calibration. Task instructions and response mappings for the run were then displayed along with a graphical reminder of the attended feature categories.

Each run comprised 256 trials, representing all possible pairs of the 16 objects, with random selection from the 100 exemplars for each class. Within each run, objects were presented in a pseudo-random order so that objects of each shape and colour were equally likely to precede objects of all shapes and colours. Each trial began with a grey fixation cross (present throughout the entire run), and the trial proceeded once fixation was verified with the eye tracker. Participants were required to fixate on the cross for at least 300ms before the stimulus would appear. Stimuli were then presented for 150ms. Finally, participants were shown the grey fixation cross on a blank screen for a variable duration. This period comprised the participant response time (maximum of 2500ms, after which time the trial proceeded; percentage of missed trials across entire study=0.69%) and a variable inter-trial interval (500-1000ms) during which fixation was verified for the subsequent trial. A variable inter-trial interval was used to mitigate expectancy effects.

Participants only received feedback (average accuracy and reaction time (RT)) at the end of each run, with the exception of five participants who additionally received feedback after every trial. For these participants, after making a response, the fixation cross changed colour for 200ms to indicate whether their response was correct (white) or incorrect (blue). Variation in feedback scheme was included as a factor in our analyses but failed to explain any variance in the behavioural or neural data.

### 2.5. Behavioural data analysis

All analyses were conducted using R (v4.2.2) in RStudio (v2023.3.0). Statistical analyses to determine whether participant behaviour (accuracy and RT) varied across task (attend to colour or shape) and feature steps (e.g., difference in accuracy between classifying ‘weakly red’ versus ‘strongly red’ object as ‘red’), were performed by fitting linear mixed-effects models (Baayen, 2008) using the *lme4* package (v1.1-32, Bates et al., 2015) and then reducing these models to identify key effects using the *lmerTest* package (v3.1-3, Kuznetsova et al., 2017).

Our full models were fitted by maximum likelihood for accuracy (*Proportion Correct*) and mean reaction time (*Average RT*). Predictors for both models comprised task (*Task*: attend to colour or shape), feature steps (*Feature Step*: e.g., for colour task, colours one to four), and their interaction (Table 1). *Subject* was defined as a random effect in both models. We only considered task-relevant information (i.e., the colour or shape of the target, in attend colour and attend shape trials, respectively) and the analysis was collapsed across attended location (left or right of fixation). For RT, only correct and feasible (i.e., within 0.15-2.5secs) trials were analysed. In addition, both models included other factors that were not of scientific interest but might capture variance (i.e., stimulus colour set and feedback scheme).

**Table 1.**
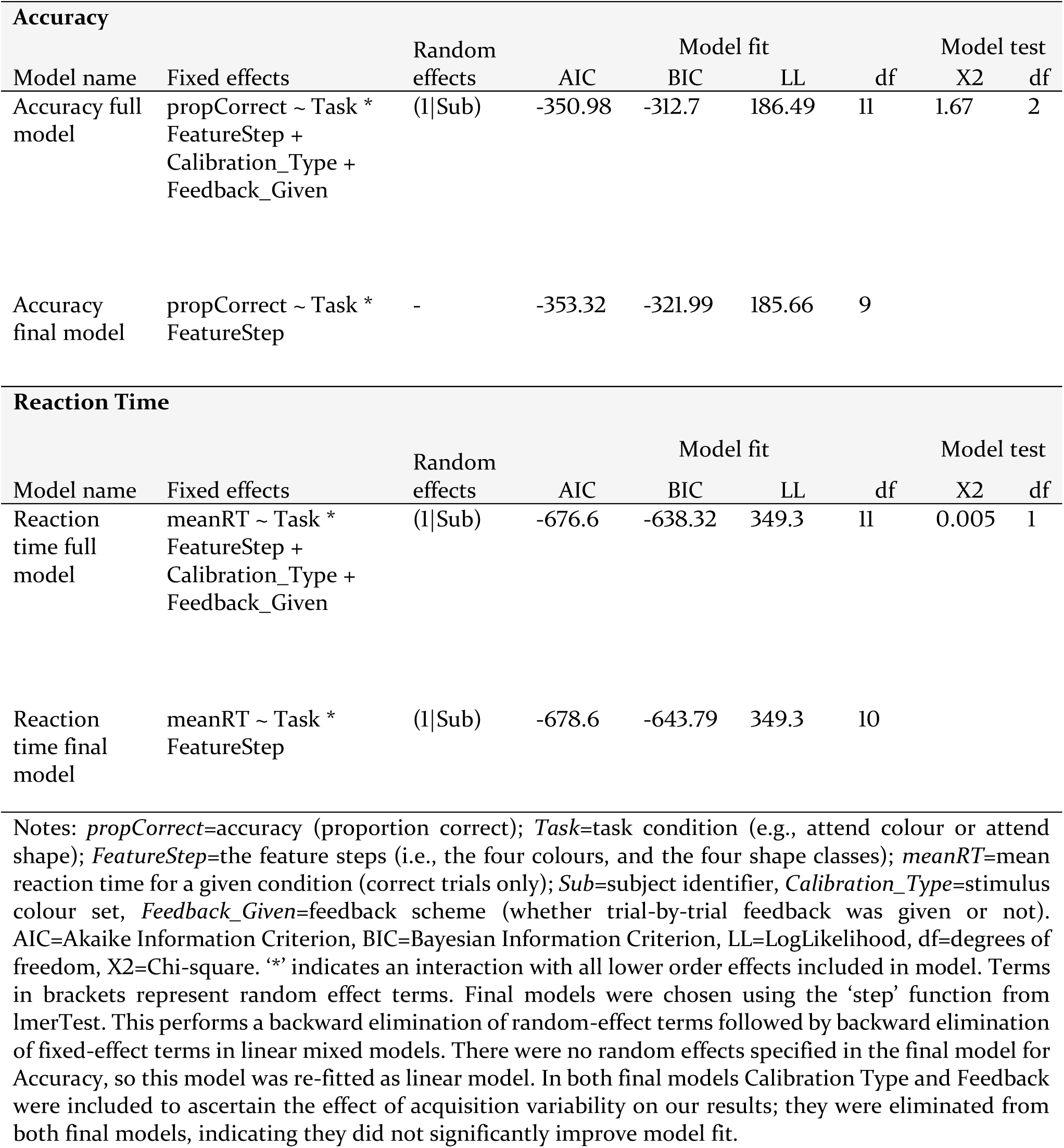
Full and final linear mixed effects models for behavioural analysis.

We used model selection to determine which fixed and random effect factors best estimated our accuracy or RT data, using the *step* function of the lmerTest package (Kuznetsova et al., 2017). This function performs a backward elimination of all effects included in our full model, starting with random effects followed by fixed effects, commencing with higher order interactions first. Our final model was that identified through this process and comprised only non-eliminated effects, namely those considered to be significant predictors of our outcome variable. Full and final models for Accuracy and RT along with model fit parameters are shown in Table 1.

These final models were re-fitted using a linear model for accuracy (as random effects were eliminated through the step procedure described earlier) and with restricted maximum likelihood for RT, to allow for significance testing of fixed effects (Meteyard & Davies, 2020). Further post-hoc tests, where necessary, were conducted using the emmeans package (v1.8.5), using the Satterthwaite method to calculate degrees of freedom where random effects were specified. P-values were adjusted for multiple comparisons using the Bonferroni method.

Extraneous variables (i.e., stimulus colour set and feedback scheme) in both accuracy and RT models were eliminated through our step model selection procedure, indicating that the inclusion of these factors did not explain significant additional variance or improve model fit (Table 1).

### 2.6. fMRI data analysis

#### 2.6.1. fMRI preprocessing

MRI data were preprocessed using SPM 12 (Wellcome Department of Imaging Neuroscience, London, UK; www.fil.ion.ucl.ac.uk/spm) in MATLAB 2019b. Functional MRI data were converted from DICOM to NIFTI format, spatially realigned to the first image, and slice time corrected. EPI images were then slightly smoothed (4mm FWHM Gaussian kernel) to improve signal-to-noise ratio, as in our previous MVPA work (e.g., Jackson et al., 2017; Woolgar et al., 2015).

Structural images were co-registered to the mean EPI, segmented into grey and white matter, and then normalised to tissue specific templates. This was done to generate individual normalisation parameters, which we used to deform template-defined regions of interest (ROIs) to native space.

#### 2.6.2. Regions of interest (ROIs)

We used the 13 frontoparietal MD ROIs from the parcellated map provided by Fedorenko et al. (2013), available online at <u>imaging.mrc-cbu.cam.ac.uk/imaging/Mdsystem</u>. These ROIs are activation-based and represent regions which demonstrate consistent recruitment across a diverse range of demanding cognitive tasks. The regions include bilateral anterior cingulate cortex (ACC; centre of mass (COM): 0 15 46.2, volume: 18.6 cm^3^), bilateral anterior insula/frontal operculum (AI/FO; COM: ±34.3 19 2.2, 7.9cm^3^), bilateral inferior frontal junction (IFJ; COM: ±43.8 3.8 32, 10.2cm^3^), bilateral anterior inferior frontal sulcus (aIFS; COM: ±34.6 47.1 18.5, 5.0cm^3^), bilateral posterior inferior frontal sulcus (pIFS; COM ±40.1 31.6 26.5, 5.7cm^3^), bilateral intraparietal sulcus (IPS; COM: ±29.4 -55.7 46.2, 34.0cm^3^), and bilateral premotor cortex (PM; COM: ±27.5 -2.3 56, 9.1cm^3^).

We also defined left and right visual cortex ROIs by taking Brodmann’s Area (BA) 17/18 (COM: -13.3, -80.6, 2.7 and 15.5 -79.2 3, respectively; volume: 56 cm^3^ and 54 cm^3^, respectively), from the Brodmann’s template of MRIcro (Rorden & Brett, 2000). Coordinates are in MNI152 space (McConnell Brain Imaging Centre, Montreal Neurological Institute, Montreal, QC, Canada).

We deformed MD and visual cortex ROIs to the native space of each participant by applying the inverse of each participant’s normalisation parameters. This allowed us to perform pattern classification analysis on an ROI basis using the native space data for each participant.

### 2.7. fMRI MVPA analysis

#### 2.7.1. General linear model (GLM)

To estimate activation patterns for MVPA, we implemented a general linear model for each participant using the realigned, slice-time-corrected, smoothed, native space EPIs with SPM12. To account for trial-by-trial differences in RT (Todd et al., 2013), trials were modelled as epochs lasting from stimulus onset until response and thus each epoch varied in length (Grinband et al., 2008; Henson, 2007; Woolgar et al., 2014), and convolved with the hemodynamic response of SPM12. We also included six movement parameters (rotation around and translation in x, y, z) and corrected for serial autocorrelation using the FAST approach (Olszowy et al., 2019).

We modelled the data with 16 regressors per run. These consisted of eight regressors (one for each of the four shapes, and one for each of the four colours) per object in the display (left and right of fixation). Each trial therefore contributed to the estimation of four regressors: the shape and colour of the object on the left, and the shape and colour of the object on the right. In combination with the attention condition (i.e., combination of attend left/right and report shape/colour), this is equivalent to modelling the target colour, target shape, distractor colour, and distractor shape on each trial. Error trials were not modelled and were excluded from the analysis.

#### 2.7.2. Stimulus information decoding

The primary aim of this study was to examine the effect of spatial and feature attention on attended and unattended stimulus information in our ROIs. To address this, we used MVPA to discriminate patterns of activation pertaining to the value (e.g., its position in colour space) of the attended and unattended feature of the object in the attended and unattended location.

We performed MVPA using the Decoding Toolbox version 3.991 (Hebart et al., 2015), which wraps the LIBSVM library (Chang & Lin, 2011). We trained separate classifiers to discriminate between each of the six possible pairings of the four feature values for shape and colour separately (e.g., colour 1 vs. colour 2, colour 1 vs. colour 3, colour 1 vs. colour 4, colour 2 vs. colour 3, colour 2 vs. colour 4, colour 3 vs. colour 4), in each of the four attention conditions. The four attention conditions were as follows: attended location attended feature (aLaF, e.g., the colour of the target object in attend colour runs), attended location unattended feature (aLuF, e.g., the colour of the target object in attend shape runs), unattended location attended feature (uLaF, e.g., the colour of the distractor object in attend colour runs), and unattended location unattended feature (uLuF, e.g., the colour of the distractor object in attend shape runs).

To illustrate how the decoding analysis proceeded, we describe the decoding analysis of colour one from colour two in the aLaF condition in detail. For a given ROI, the pattern of beta values was extracted from each of the two relevant beta images (one for each colour) in each of the four relevant runs (four attend colour runs, comprising two attend left and two attend right runs, and both possible response mappings), yielding eight multivoxel vectors. All the voxels in each ROI contributed to each vector, without feature selection. We used a linear support vector machine (LibSVMC, cost parameter C=1) to classify between the four vectors pertaining to colour one (‘strongly red’) and the four vectors pertaining to colour two (‘weakly red’). As the four vectors comprising each classification included both stimulus locations, the classifier generalised across stimulus location. We split the data into training and testing chunks using leave-one-run-out cross-validation and iterated around all combinations so that all runs contributed equally to training and testing. The four classification accuracies from the four cross-validation folds were then averaged to give a single accuracy score for that participant, ROI, attention condition, and pair of colours. We repeated this procedure for each pairwise comparison and each of the four attention conditions separately. For all decoding analyses, we use percentage correct to express classifier accuracy, where chance performance has a theoretical value of 50%.

#### 2.7.3. Model selection and statistical analyses

As with our behavioural analyses, statistical analyses to examine the effect of spatial and feature attention on stimulus information coding, across ROIs and across stimulus space, were performed by fitting linear mixed-effects models using the *lme4* package (v1.1-32, Bates et al., 2015) and then reducing these models to identify key effects using the *lmerTest* package (v3.1-3, Kuznetsova et al., 2017) in R.

Our full models comprised all fixed and random effects of theoretical importance (see Table 2 for details of our full and final models). In all cases, models were fit to the accuracy of the classification (i.e., strength of information coding) of the given feature being decoded (*Feature Decoded*: colour/shape, averaged across all pairwise classifications), across each of the different attention conditions (*Location Attended * Feature Attended*: aLaF, aLuF, uLaF, uLuF). We then tailored our full models to answer different research questions by specifying additional fixed effects as appropriate. Our first research question examined whether spatial and feature attention modulated stimulus information coding; we examined these attention effects in MD and visual regions in two separate models (an MD model and a visual region model). For the MD model we included *ROIs* as a predictor in our full MD model. For both MD and visual region models, we also included variables that might capture additional variance (i.e., stimulus colour set and feedback scheme). Our second research question examined whether the effect of attention varied across stimulus space. We again analysed this in separate models for MD and visual regions. We included *Distance in Feature Space* as a predictor in both models. This predictor reflected the physical discriminability of the stimuli with three levels: large (pairwise comparisons of three feature steps: one and four), medium (pairwise comparisons of two feature steps: one and three, two and four), and small (pairwise comparisons of one feature step: one and two, two and three, three and four). All predictors were included as fixed effects with all possible interaction terms included. As we reasoned that classification accuracy may vary across subjects as a function of attention condition, attention condition and subject were defined as random effects. As we sought to compare models primarily on the basis of fixed effects, we fit the full models using maximum likelihood as is recommended in the literature (e.g., see Meteyard & Davies, 2020). We used model selection to determine which fixed and random effect factors best estimated decoding accuracy using the *step* function of the lmerTest package (Kuznetsova et al., 2017). The final models comprised effects that were identified as significant predictors of classification accuracy. As in the behavioural analysis, final models were re-estimated with restricted maximum likelihood to allow for significance testing of fixed effects (Meteyard & Davies, 2020). Post-hoc analyses were performed using the emmeans package (v 1.8.5), using the Satterthwaite method to calculate degrees of freedom. We used Bonferroni adjustment to correct for multiple comparisons.

**Table 2.** Full and final models specified for decoding analysis for MD and visual regions and final model selected.

| How do spatial and feature attention modulate the coding of stimulus information? |  |  |  |  |  |  |  |  |
| --- | --- | --- | --- | --- | --- | --- | --- | --- |
| MD regions |  |  |  |  |  |  |  |  |
| Model name | Fixed effects | Random effects | AIC | Model fit |  | df | Model test |  |
|  |  |  |  | BIC | LL |  | X2 | df |
| Full model | Decoding Accuracy ~ Location Attended * Feature Attended * Feature Decoded * ROIs + CalibrationType + feedbackGiven | (Location Attended * Feature Attended \| Sub) | 88096 | 88594 | -43979 | 69 | 12.93 | 26 |
| Final model | Decoding Accuracy ~ Location Attended + Feature Attended + Feature Decoded + ROIs + Location Attended:Feature Attended + Location Attended: Feature Decoded + Feature Attended: Feature Decoded + Location Attended:ROIs + Feature Attended:ROIs + Location Attended:Feature Attended: Feature Decoded + Location Attended:Feature Attended:ROIs | (Location Attended * Feature Attended \| Sub) | 88057 | 88367 | -43986 | 43 |  |  |

How do spatial and feature attention modulate the coding of stimulus information? Visual regions
| Model name | Fixed effects | Random effects | Model fit |  |  | Model test |  |  |
| --- | --- | --- | --- | --- | --- | --- | --- | --- |
|  |  |  | AIC | BIC | LL | df | X2 | df |
| Full model | Decoding Accuracy ~<br>Location Attended *<br>Feature Attended *<br>Feature Decoded +<br>CalibrationType +<br>feedbackGiven | (Location Attended *<br>Feature Attended<br>\| Sub) | 12747 | 12857 | -6352.3 | 21 | 4.03 | 4 |
| Final model | Decoding Accuracy ~<br>Location Attended +<br>Feature Attended +<br>Feature Decoded +<br>Location Attended:Feature<br>Attended + Feature<br>Attended: Feature<br>Decoded | (Location Attended *<br>Feature Attended<br>\| Sub) | 12743 | 12832 | -6354.4 | 17 |  |  |

Does the effect of selective attention vary across stimulus space? MD regions
| Model name | Fixed effects | Random effects | Model fit |  |  | Model test |  |  |
| --- | --- | --- | --- | --- | --- | --- | --- | --- |
|  |  |  | AIC | BIC | LL | df | X2 | df |
| Full model | Decoding Accuracy ~<br>Location Attended *<br>Feature Attended *<br>Distance in feature space *<br>Feature Decoded | (Location Attended *<br>Feature Attended<br>\| Sub) | 87899 | 88152 | -43915 | 35 | N/A |  |
| Final model | Final model=full model |  |  |  |  |  |  |  |

Does the effect of selective attention vary across stimulus space? Visual regions
| Model name | Fixed effects | Random effects | Model fit |  |  |  | Model test |  |
| --- | --- | --- | --- | --- | --- | --- | --- | --- |
|  |  |  | AIC | BIC | LL | df | X2 | df |
| Full model | Decoding Accuracy ~<br>Location Attended *<br>Feature Attended *<br>Distance in feature space *<br>Feature Decoded | (Location Attended<br>*<br>Feature Attended<br>\| Sub) | 12749 | 12933 | -6339.4 | 35 | 29.93 | 18 |
| Final model | Decoding Accuracy ~<br>Location Attended +<br>Feature Attended +<br>Feature Decoded +<br>Location Attended:Feature Attended +<br>Feature Attended: Feature Decoded | (Location Attended<br>* Feature Attended<br>\| Sub) | 12743 | 12832 | -6354.4 | 17 |  |  |

Notes: *Decoding Accuracy*=average decoding accuracy for given comparison, *Location Attended*=binary variable indicating whether decoding comparison was for the object in the attended location, *Feature Attended*=binary variable indicating whether decoding comparison was for the attended feature, *Feature Decoded*=binary variable indicating whether decoding of colour or shape information, *ROIs*=each of the MD ROIs included in this analysis, *Distance in Feature Space*=the physical discriminability of the stimuli with three levels: large (pairwise comparisons of three feature steps: one and four), medium (pairwise comparisons of two feature steps: one and three, two and four), and small (pairwise comparisons of one feature step: one and two, two and three, three and four), *Sub*=subject identifier, *Calibration\_Type*=stimulus colour set, *Feedback\_Given*=whether trial-by-trial feedback was given or not. AIC=Akaike Information Criterion, BIC=Bayesian Information Criterion, LL=LogLikelihood, df=degrees of freedom, X2=Chi-square. Note: ‘.’ indicates interaction between only the terms specified (i.e., without lower-order effects), while ‘\*’ indicates an interaction with all lower order effects included in model. Terms in brackets represent random effect terms. Final models were chosen using the ‘step’ function from lmerTest. This performs a backward elimination of random-effect terms followed by backward elimination of fixed-effect terms in linear mixed models.

Table 2 shows the full (i.e., initially specified) and final models, along with model fit parameters, in MD and visual regions. As with the behavioural results, our step model selection procedure eliminated extraneous variables (i.e., stimulus colour set and feedback scheme) from our final models, indicating that the inclusion of these factors did not significantly explain any additional variance or improve model fit.

### 2.8. Generalisation analyses

As a secondary analysis, we probed the organisational structure of task-related representations in MD regions. For this, we ran a series of cross-classification decoding analyses using only task-relevant (i.e., aLaF) information.

First, we examined whether we could decode ‘difficulty’ (i.e., ‘easy’ from ‘hard’ stimuli) from our data. To illustrate the procedure, we describe the process for colour information. We first trained our classifier to distinguish between two features on the same side of the decision boundary that reflected two levels of difficulty (e.g., ‘reddish’: colour one (strongly red, *easy*) vs colour two (weakly red, *hard*)). We then tested this classifier on whether it would correctly classify the difficulty of the two features on the other side of the decision boundary (e.g., ‘greenish’: colour four (strongly green, *easy*) vs colour three (weakly green, *hard*)). This procedure was repeated in the reverse (i.e., training to distinguish between easy green and hard green, and testing on easy red and hard red). This analysis allowed us to examine whether a representation of ‘difficulty’, as defined by distinguishing between hard and easy trials on one side of the decision boundary, generalised across feature space. We then performed the equivalent analysis for shape information.

Second, we tested whether we could decode the feature categories (i.e., for colour: ‘reddish’ or ‘greenish’, for shape: ‘X-shaped’ or ‘non-X-shaped’), independent of difficulty level. We first trained our classifier to distinguish between two features on opposite sides of the decision boundary but equidistant from it, that is, matched in terms of difficulty (e.g., colour one (strongly *red*, easy) and colour four (strongly *green*, easy)). We then tested whether this classifier would correctly classify the colour of the remaining two features (e.g., colour two (weakly *red*, hard) and colour three (weakly *green*, hard)), and vice versa. We then repeated the same procedure for shape information. This analysis allowed us to examine whether the distinction between feature categories (e.g., red and green), generalised across difficulty (i.e., easy and hard).

### 2.9. Comparison to chance

Many of the effects in our statistical models would only be interpretable if decoding in one or more of the conditions was also significantly above chance. In these cases, we used the Bayes Factor package in R (Morey & Rouder, 2018) to quantify evidence for above-chance (H1) or at-chance (null) classification accuracy. We defined priors using a half-Cauchy prior for the alternative, with a default width of 0.707 and an interval ranging from a standardised effect size of 0.5 to infinity (Jeffreys, 1998; Rouder et al., 2009; Wetzels et al., 2011), as in our previous work (Moerel et al., 2021; Teichmann et al., 2021). We used a point null centred on chance (Morey & Rouder, 2011). We report the Bayes Factor (BF10), which indicates the likelihood that the evidence favours the alternative hypothesis relative to the null hypothesis. For instance, a BF10 of 3 indicates that the data were three times more likely to occur under the alternative than the null hypothesis. BF10s > 3, 10, 30, and 100 are considered ‘moderate’, ‘strong’, ‘very strong’, and ‘extreme’ evidence in support of the alternative hypothesis (i.e., decoding accuracy above chance), respectively (Wagenmakers et al., 2018). Conversely, BF10s < 1/3, 1/10, 1/30 and 1/100 are considered ‘moderate’, ‘strong’, ‘very strong’, and ‘extreme’ evidence in support of the null hypothesis (i.e., at-chance decoding accuracy).

### 2.10. Searchlight analysis

To examine the extent to which the multiplicative effect was specific to the MD and visual cortex, and whether evidence of irrelevant information coding could be found in other brain regions, we ran whole-brain searchlight analyses (Kriegeskorte et al., 2006). Since the data used in a searchlight analysis overlaps with that of the ROIs, we would expect some convergence between the two results. However, the benefit of a searchlight is that it allows us to determine whether any additional regions show similar effects when looking at information coding on a finer spatial scale.

For each participant, data were iteratively extracted from a sphere centred on each voxel in the brain of radius 3 voxels (∼10mm), as in our previous work (e.g., Jackson & Woolgar, 2018). A linear support vector machine was trained and tested as for our ROI analysis above (Section 2.7), but using data from each sphere. The classification accuracy value for that sphere was then assigned to the central voxel. This yielded whole-brain accuracy maps for each individual, for each attention condition. To allow for group level analysis, accuracy maps were normalised using normalisation parameters extracted at pre-processing stage, and smoothed using an 8 mm full-width at half-maximum (FWHM) Gaussian kernel. Finally a one-sample t-test was conducted at each voxel to identify where classification was significantly above chance. The results were family-wise error (FWE) corrected at p=0.0000001, with an extent threshold of 20 voxels. All coordinates are given in MNI152 space (McConnell Brain Imaging Centre, Montreal Neurological Institute, Montreal, QC, Canada).

## 3. Results

### 3.1. Behavioural results

The final LME models selected through our model selection process for accuracy and RT are shown in Table 1. There was a significant effect of feature step on both RT and accuracy (main effect of feature step: RT: F(3,203)=118.95, p<0.0001, accuracy: F(3,232)=87.87, p<0.0001, see Supplementary Figure S 1). This main effect was modulated by a significant interaction with task, (task and feature step interaction for RT: F(3,203)=19.68, p<0.0001; accuracy: F(3,232)=21.25, p<0.0001), but post-hoc tests for each task showed that the main effect of feature step was significant for both tasks separately (corrected p-values < 0.05 for colour and shape). More specifically, post-hoc tests showed that for the shape task, participants were faster and more accurate when classifying easy (shape 1 and 4) than hard (shapes 2 and 3, all p values < 0.05) shapes. Similarly, for the colour task, participants were faster and more accurate in classifying easy (colours 1 and 4) than hard (colours 2 and 3, all p-values <0.05) colours. However, they were also significantly less accurate and slower in classifying colour two (weakly red) than any other colour (all p-values <0.0001). Thus, as expected, participant behaviour reflected a tendency to be faster and more accurate on the easy compared to the hard stimuli.

Participants were also significantly faster to identify colours than shapes (colour mean RT=0.71±0.13s, shape mean RT=0.72±0.10s; main effect of task on RT: F(1,210)=7.20, p=0.008) but there was no evidence that they differed in accuracy between the two tasks (colour mean accuracy=0.84±0.21, shape mean accuracy=0.86±0.13; main effect of task on accuracy: F(1,232)=2.31, p=0.130).

### 3.2. fMRI MVPA results: Stimulus information coding

#### 3.2.1. MD regions selectively code of task-relevant information

We next turned to the fMRI data to consider our key question of whether spatial and feature attention modulated the coding of stimulus information and, if so, whether the two subtypes of attention interacted in modulating the neural code. Figure 2 shows the effect of these two subtypes of attention on decoding of stimulus information. We found that, indeed, both spatial attention (F(1,29)=90.13, p<0.0001) and feature attention (F(1,29)=59.09, p<0.0001) boosted stimulus information processing, and there was a significant two-way interaction between the two subtypes of attention F(1,29)=67.59, p<0.0001). As shown in Figure 2 (Panel A), the multiplicative interaction resulted in information being decodable only if a stimulus feature was both relevant to the task and at the focus of spatial attention (BF10>100 for the attended feature information at the attended location, i.e., aLaF, with BF10s<0.1 for other attention conditions).

**Figure 2.**
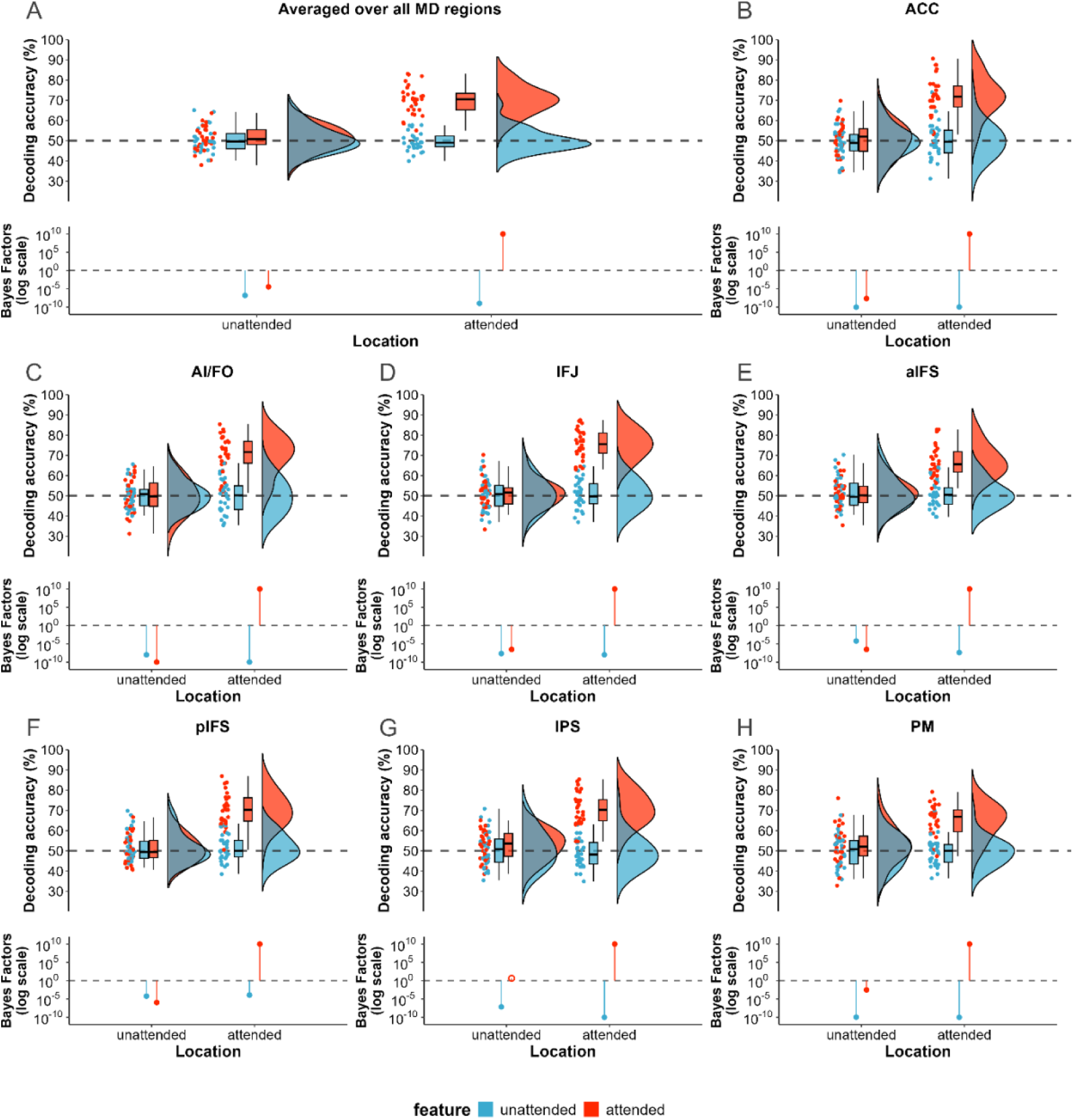
The effect of two interacting subtypes of attention on decoding of feature information. Decoding accuracy reflects average accuracy in decoding colour or shape information in the object that was in the unattended (left plots) or attended (right plots) location, when the feature in question was attended (red) or unattended (blue). Dots depict individual subject results, boxplots indicate median (central line), first and third quartiles (box hinges), and largest and smallest decoding accuracy value within 1.5 x applicable quartile range (whiskers), half violin depicts data distribution. The strength of evidence for above-chance (H1) or at-chance (H0) decoding was quantified using Bayesian t-tests and is shown as Bayes Factors for each of the conditions on a logarithmic scale, with solid circles reflecting at least moderate evidence in favour of H1 or H0, and open circles reflecting evidence was anecdotal or inconclusive. Panel A: average results across all MD ROIs. Panels B-H: each MD region, shown separately. B: anterior cingulate cortex (ACC); C: anterior insula/frontal operculum (AIFO); D: inferior frontal junction (IFJ); E: anterior inferior frontal sulcus (aIFS); F: posterior inferior frontal sulcus (pIFS); G: intraparietal sulcus (IPS); H: premotor cortex (PM).

The key two-way interaction was modulated by two three-way interactions that did not change the interpretation. First, there was a three-way interaction between spatial attention, feature attention and MD region (F(6,9932)=3.51, p=0.002). This reflected variation in the strength of the two-way interaction over MD regions with the most prominent effects in IFJ and ACC and smallest effects in PM (see Figure 2, panels B:H). Nevertheless, the multiplicative interaction was significant in all regions separately (all p-values<=0.0001), with selective boosting of only attended feature information at the attended location across all regions (all BF10s>100 for aLaF information; anecdotal evidence for above-chance decoding of uLaF in the IPS; else BF10s<0.3 for other attention conditions across all MD regions).

Second, there was a three-way interaction between spatial attention, feature attention and feature decoded (F(1,9932)=4.81, p=0.028) (see Supplementary Figure S 2). This reflected a larger multiplicative effect of the two types of attention on coding of object colour, relative to shape. Nevertheless, the two-way interaction was significant for each feature separately (colour: F(1,35.18)=72.24, p<0.001; shape: F(1,35.18=51.40, p<0.001), and extreme evidence for above-chance decoding was found for only the attended feature of the target object (BF10s>100; BF10s<0.38 for all other attention conditions).

#### 3.2.2. Task-relevant information is organised across two dimensions in MD regions

The results from our first analysis indicate a multiplicative effect of spatial and feature attention on representations in MD regions such that only information about the attended feature of the target object was enhanced. In previous work with similar stimuli, we found that spatial and feature-selective attention had qualitatively different effects on coding of stimulus information, with spatial attention tending to emphasise large feature differences, and feature-selective attention tending to enhance small feature differences (Goddard et al., 2022), in line with Reynolds and Heeger’s (2009) normalisation model. Therefore, as a secondary analysis, here we asked whether the selective boost in the representation of task-relevant information was uniform across stimulus space.

Our results suggested that the interacting effect of spatial and feature attention on information coding varied with the physical discriminability of the stimuli (three-way interaction spatial attention * feature attention * distance in feature space, F(2,9940)=31.07, p<0.0001, Figure 3). This effect was more pronounced for colour than shape information (four-way interaction, F(2,9940)=6.03, p=0.002), but significant for both colour and shape information separately (colour: F(2,9940) p<0.0001)=31.36; shape: F(2,9940)=5.74, p=0.003).

**Figure 3.**
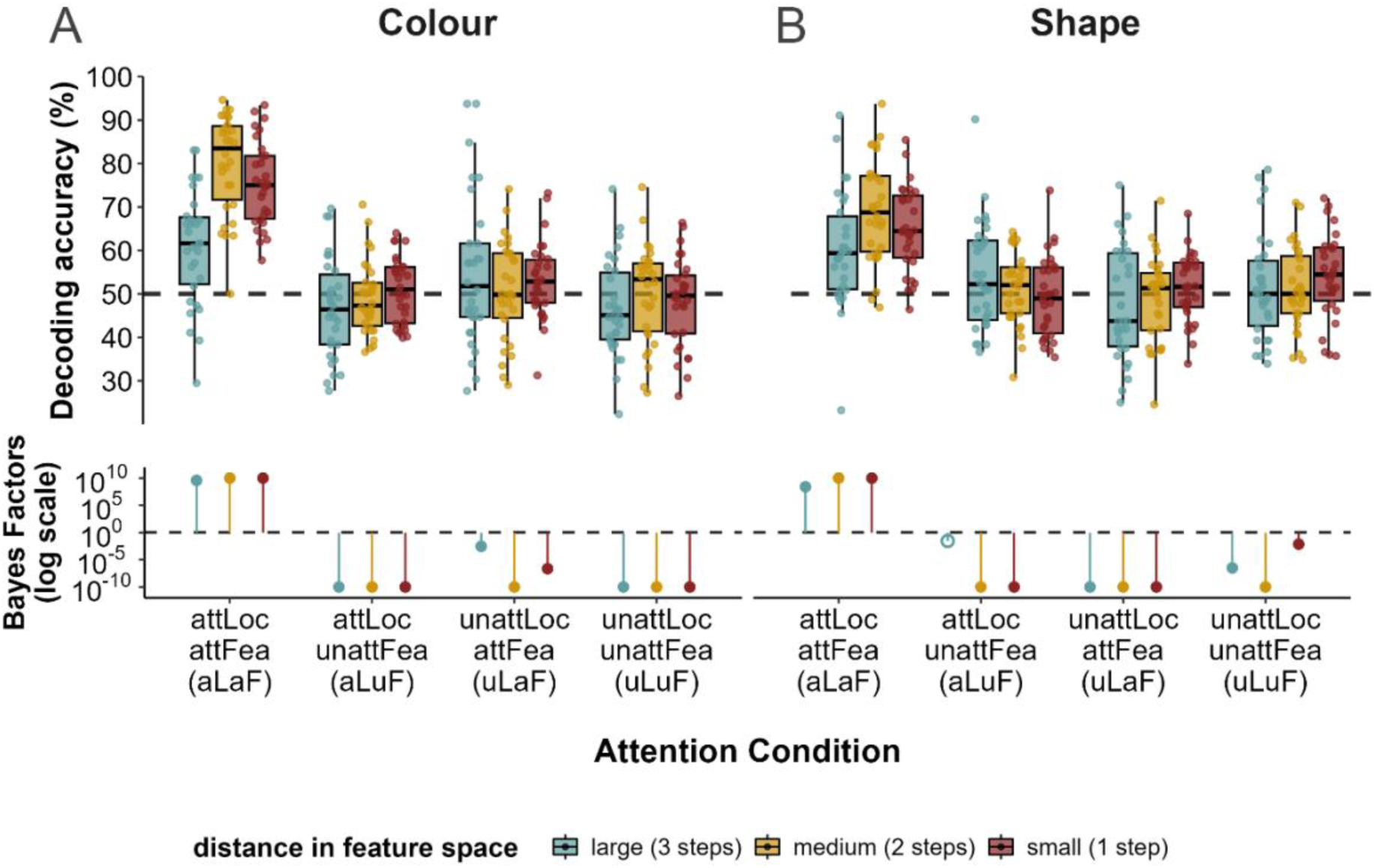
Decoding of colour (Panel A) and shape (Panel B) information in each of the spatial and feature attention condition for each distance in feature space, collapsed across all MD regions. Distance in feature space levels are large (pairwise comparisons of three feature steps: 1v4), medium (pairwise comparisons of two feature steps: 1v3, 2v4), and small (pairwise comparisons of one feature step: 1v2, 2v3, 3v4). Dots depict individual subject results, boxplots indicate median (central line), first and third quartiles (box hinges), and largest and smallest decoding accuracy value within 1.5 x applicable quartile range (whiskers). The strength of evidence for above-chance (H1) or at-chance (H0) decoding was quantified using Bayesian t-tests and is shown as Bayes Factors for each of the conditions on a logarithmic scale, with solid circles reflecting at least moderate evidence in favour of H1 or H0, and open circles reflecting evidence was anecdotal or inconclusive.

The differential attention effect over step size appeared to be driven by differences in the coding of task-relevant (aLaF) information as all other conditions were at chance (mean decoding accuracy 60.2-80.0% and BFs>100 for aLaF conditions; for all other attention conditions, mean decoding accuracy 46.7-55.0%; all BF10s<0.34; Figure 3). Within the aLaF condition, post-hoc tests revealed that medium (two step) physical discriminations were more readily decoded than small (one step) (colour: t(9940)=3.90, p<0.001; shape: t(9940)=3.63, p<0.001), or large (three step) (colour: t(9940)=12.11, p<0.0001; shape: t(9940)=5.77, p<0.0001) discriminations. In addition, small physical discriminations were decoded more readily that large ones (colour: t(9940)=9.77, p<0.0001); shape: t(9940)=3.25, p=0.004). This suggested that attention boosted the representation of task-relevant information in MD regions differentially according to the physical discriminability of the stimuli, with advantage for medium and then small physical discriminations over large ones.

In this analysis, medium and large discriminations always crossed the stimulus decision boundary used by the participant, while a subset of the ‘small’ discriminations (step sizes one and two, and step sizes three and four) did not. To check whether this difference contributed to our results, we re-ran the analysis including only the pairs that straddled the decision boundary (i.e., only the step size two and three pairwise comparison was included for the small step size). This did not change the overall pattern of results: the small step size was now as decodable as the large step size, but the medium step size was still more decodable than small and large step sizes (Supplementary Figure S 3).

Since the medium step size classification always discriminated between one perceptually easier and one perceptually harder item, it was possible that medium step size decoding reflected sensitivity to difficulty differences as well as feature differences between the stimuli. To explore this, we asked to what extent difficulty was an organising dimension for the MD representations in this task. We found that coding of item difficulty (easy vs hard) generalised over each half of feature space (e.g., from red to green and vice versa) for both colour (Figure 4, Panel A) and shape (Figure 4, Panel B). There was strong to extreme evidence for this effect across all MD regions (colour mean accuracy range=64.2-73.6%, all BF10s>100; shape mean accuracy range=58.2-67.7%, all BF10s>46.80), with the exception of shape information in IPS, where the evidence of stimulus coding was anecdotal (mean accuracy=57.0%, BF10=1.82).

**Figure 4.**
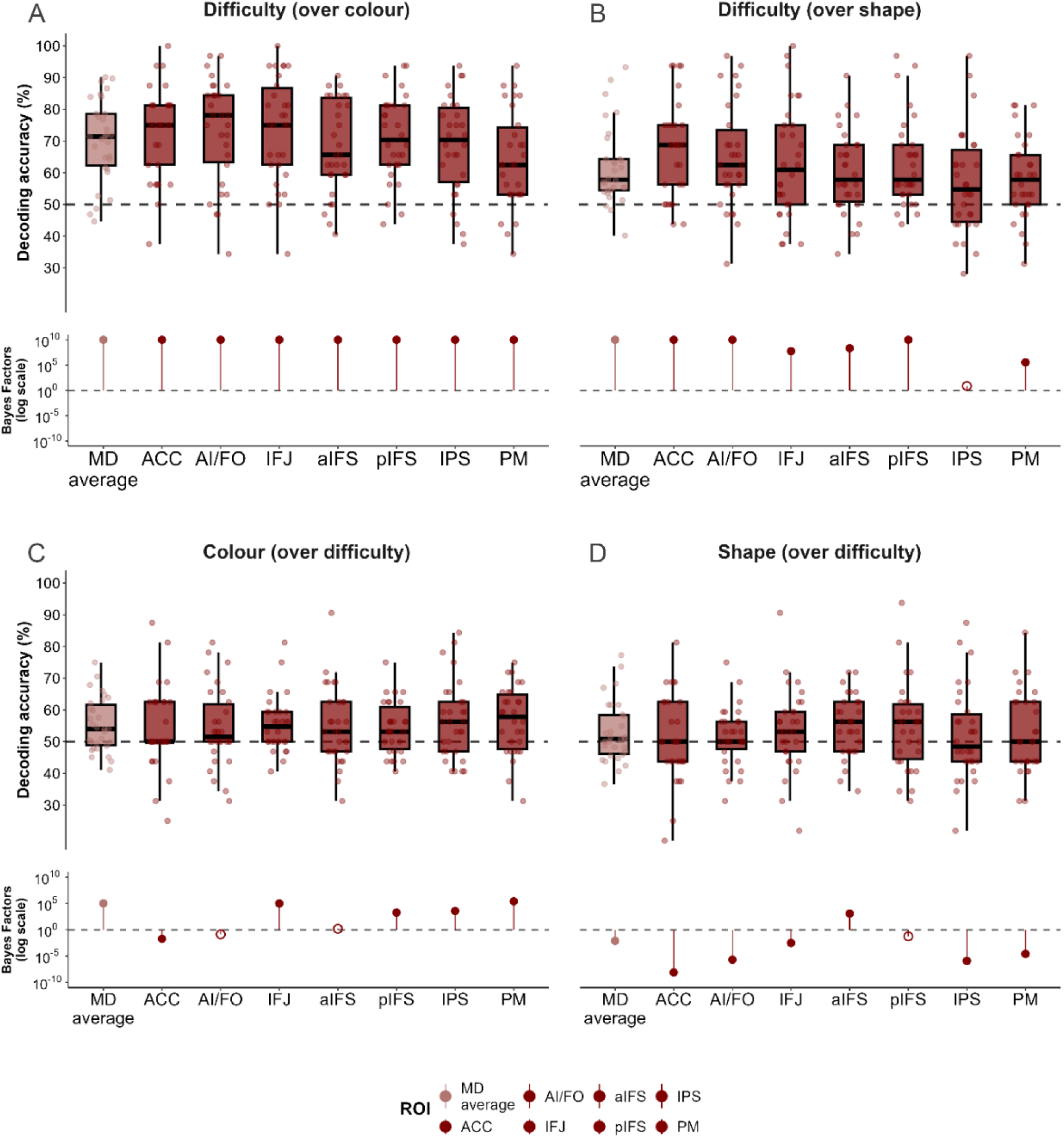
Top panels: decoding of difficulty generalised over the two colour categories (Panel A) and shape categories (Panel B). Bottom panels: decoding of colour category information (Panel C) and shape category information (Panel D) generalised over difficulty. Decoding accuracy shown for each MD region (collapsed across hemisphere, light red), and averaged across MD ROIs (dark red). For task-relevant information (i.e., aLaF) only. Dots depict individual subject results, boxplots indicate median (central line), first and third quartiles (box hinges), and largest and smallest decoding accuracy value within 1.5 x applicable quartile range (whiskers). The strength of evidence for above-chance (H1) or at-chance (H0) decoding was quantified using Bayesian t-tests and is shown as Bayes Factors for each of the conditions on a logarithmic scale, with solid circles reflecting at least moderate evidence in favour of H1 or H0, and open circles reflecting evidence was anecdotal or inconclusive.

Next, we analysed the reverse: whether representation of stimulus category (for colour: red or green, for shape: X-shaped or non-X-shaped), generalised over difficulty (from easy to hard and vice versa). We found very strong evidence that coding of colour generalised over difficulty level for MD regions overall (lighter bar in Figure 4, Panel C: mean accuracy=55.2%, BF10=32.31), and strong evidence for this effect in the IPS, pIFS, IFJ and PM regions separately (accuracy range=54.4-56.7%, BF10s>9.71). In contrast, we found moderate evidence for the null in the equivalent analysis for shape information in MD regions overall (lighter bar in Figure 4, Panel D: mean accuracy=52.9%, BF10=0.23), with only aIFS showing moderate evidence that coding of shape generalised over difficulty (mean accuracy=55.1%, BF10=8.48). These results suggest that responses in MD regions reflect both task-relevant stimulus categorisations (especially for colour, e.g., red vs green) and the difficulty of those discriminations (easy vs hard).

#### 3.2.3. Selective prioritisation of task-relevant information extends to early visual cortex

Next, we sought to contextualise the attentional effects seen in MD regions by examining the effects of attention on stimulus processing elsewhere in the brain. To look at this, we ran the same set of analyses in early visual cortex as an ROI (BA17/18), and on a whole brain basis using a roaming searchlight.

In visual cortex, we first asked whether spatial and feature attention modulated the coding of stimulus information, and if so, whether these two subtypes of attention interacted in a multiplicative manner, as we had seen in MD regions. As shown in Table 2, there were significant main effects of spatial attention (F(1,29)=16.21, p<0.001) and feature attention (F(1,29)=14.00, p<0.0001) that were modulated by a significant two-way interaction between spatial and feature attention (F(1,29)=16.73, p<0.001), that did not differ between colour and shape information (see final model in Table 2). As we had observed in MD regions, information coding was strongest for the attended feature of the target object (Figure 5), with only this task-relevant information decodable above chance (BF10>100 for aLaF; BF10s for all other attention conditions <0.1).

**Figure 5.**
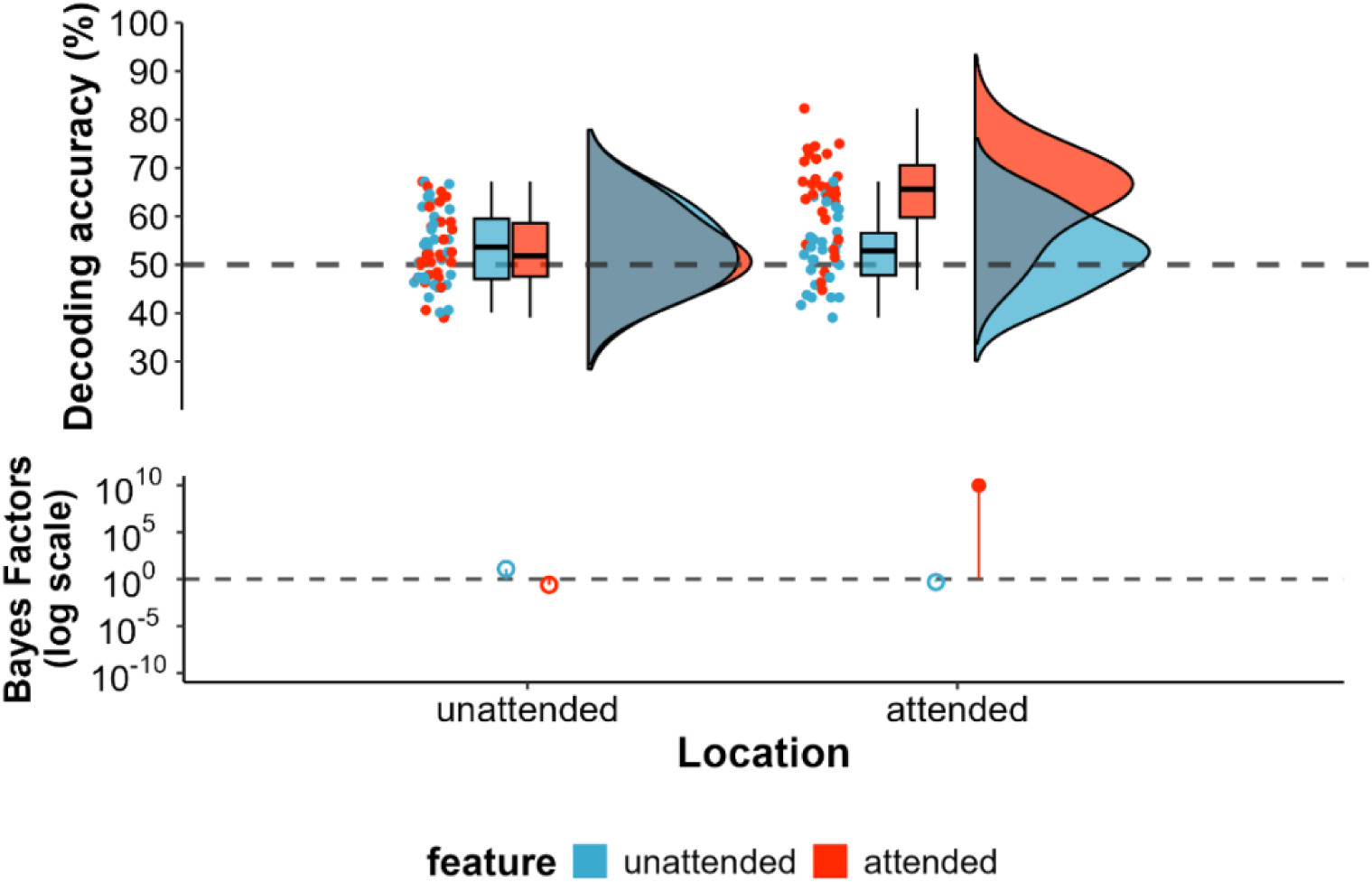
The effect of two interacting subtypes of attention on decoding of feature information for visual cortex (BA17/18). Decoding accuracy reflects average accuracy in decoding colour or shape information in the object that was in the unattended (left plots) or attended (right plots) location, when the feature in question was attended (red) or unattended (blue). Dots depict individual subject results, boxplots indicate median (central line), first and third quartiles (box hinges), and largest and smallest decoding accuracy value within 1.5 x applicable quartile range (whiskers), half violin depicts data distribution. The strength of evidence for above-chance (H1) or at-chance (H0) decoding was quantified using Bayesian t-tests and is shown as Bayes Factors for each of the conditions on a logarithmic scale, with solid circles reflecting at least moderate evidence in favour of H1 or H0, and open circles reflecting evidence was anecdotal or inconclusive.

#### 3.2.4. Attention effects do not differ across feature space in visual cortex

Next, we asked whether these selective attention effects varied across stimulus space in visual cortex. We anticipated that bottom-up factors (physical discriminability) might lead to a stronger representation of larger feature differences (i.e., greater decoding of features further apart in feature space) than smaller differences in visual cortex. On the other hand, top-down effects (e.g., feedback), might tend to drive representations towards the strong discrimination of medium step sizes, as seen in MD regions.

In fact, although the pattern of effects was visually similar to those of MD regions (Figure 6), there was no statistical evidence that attention effects varied across stimulus space for visual cortex. Table 2 shows the full model initially specified in our linear mixed-effects analysis and the final model selected through our analysis. Inclusion of stimulus discriminability (or distance in feature space) did not explain any additional variance above and beyond that explained by spatial attention, feature attention, colour or shape information, or their interactions (compare full and final models specified in Table 2; distance in feature space was not included in final model). As for the MD analysis, we also tested the same effect after restricting the analysis to only include comparisons that straddle the decision boundary; the results of this modified analysis were broadly the same, with no effect of stimulus discriminability on the spatial and feature interaction (see Supplementary Figure S 4). Overall, the qualitative pattern of the effect across feature space was similar, but weaker, than in MD regions, perhaps reflecting a combination of bottom-up and top-down factors on coding in this region.

**Figure 6.**
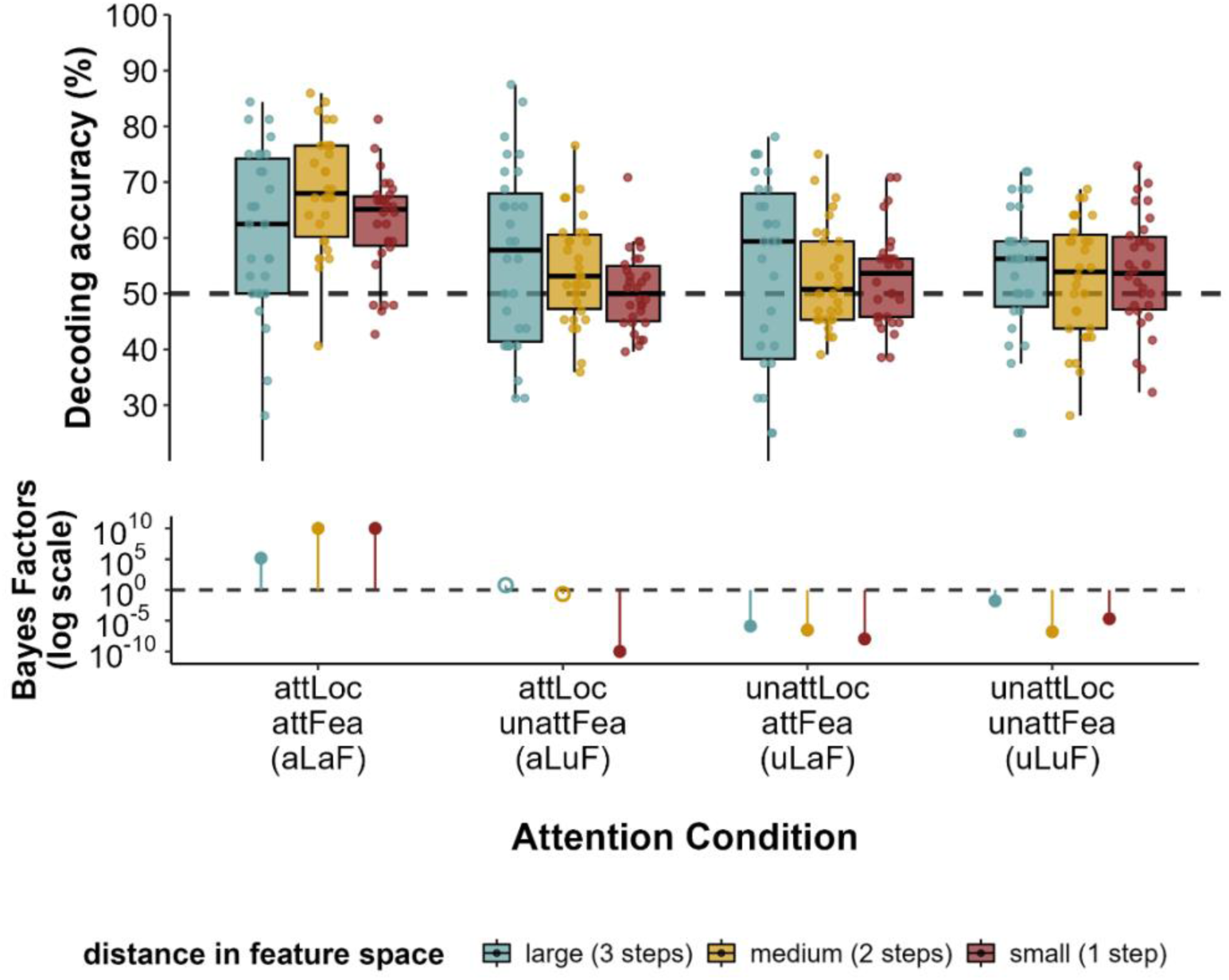
Decoding of stimulus information (averaged over colour and shape information, as not significantly different between these types of information) for visual cortex (BA17/18). Distance in feature space levels are large (pairwise comparisons of three feature steps: 1v4), medium (pairwise comparisons of two feature steps: 1v3, 2v4), and small (pairwise comparisons of one feature step: 1v2, 2v3, 3v4). Dots depict individual subject results, boxplots indicate median (central line), first and third quartiles (box hinges), and largest and smallest decoding accuracy value within 1.5 x applicable quartile range (whiskers). The strength of evidence for above-chance (H1) or at-chance (H0) decoding was quantified using Bayesian t-tests and is shown as Bayes Factors for each of the conditions on a logarithmic scale, with solid circles reflecting at least moderate evidence in favour of H1 or H0, and open circles reflecting evidence was anecdotal or inconclusive.

#### 3.2.5. Searchlight results

Finally, we ran a whole brain searchlight to characterise the spatial distribution of feature information. For task-relevant information, the searchlight results indicated widespread representation throughout the brain. To visualise the peaks of the decoding map (Figure 7), we used a cluster-level family wise error (FWE) correction for multiple comparisons with a high threshold (p<0.000001). The surviving clusters showed reasonable overlap with frontoparietal MD regions (see Figure 7 and Table 3). There was an additional prominent cluster in mid temporal cortex, particularly on the left. Intriguingly, more recent anatomical definitions of the MD system include a temporal MD region close to this cluster (Assem, Glasser, et al., 2020).

**Figure 7.**
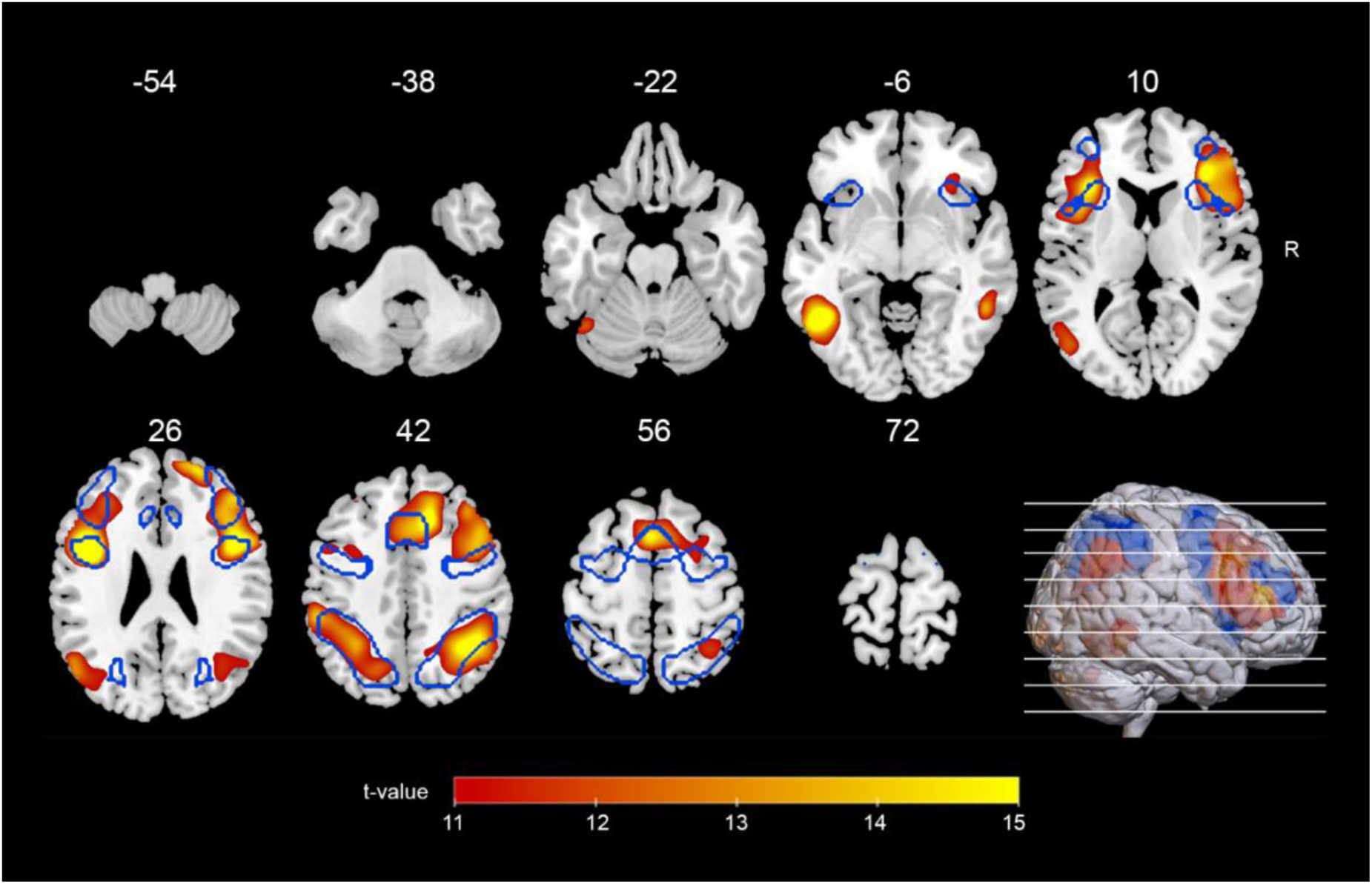
Searchlight decoding results for task-relevant (aLaF) stimulus information only. Outline of MD regions from Fedorenko (2013) are overlaid in blue. Results are thresholded at t=11.28, equivalent to p<0.0000001, FWE corrected. No stimulus information was decodable for other attention conditions at this threshold.

**Table 3.** Searchlight results.

| Attention condition | Cluster | Hemisphere | Coordinates |  |  | t-score | Cluster Extent |
| --- | --- | --- | --- | --- | --- | --- | --- |
|  |  |  | x | y | z |  |  |
| aLaF | inferior temporal gyrus, middle temporal gyrus, fusiform gyrus | Left | -50 | -54 | -10 | 20.54 | 989 |
|  | inferior frontal gyrus, middle frontal gyrus, precentral gyrus, insula | Left | -44 | 6 | 26 | 19.09 | 3582 |
|  | superior frontal gyrus (r), middle frontal gyrus (r), inferior frontal gyrus (r), precentral gyrus (r), insula (r) , supplementary motor area (r and l) , cingulum (r and l), superior frontal gyrus (l) | Right peak, extending into left | 22 | 56 | 22 | 17.56 | 7442 |
|  | inferior parietal, angular gyrus, supramarginal gyrus, middle occipital gyrus | Right | 36 | -52 | 46 | 16.18 | 2272 |
|  | middle temporal gyrus, angular gyrus, middle occipital gyrus | Left | -50 | -66 | 20 | 14.23 | 696 |
|  | inferior parietal , superior parietal, angular gyrus, supramarginal gyrus, middle occipital gyrus, superior occipital gyrus | Left | -54 | -34 | 40 | 14.08 | 1532 |
| aLuF | inferior temporal gyrus, middle temporal gyrus | Right | 58 | -62 | -2 | 7.67 | 69 |
|  | inferior occipital | Left | -36 | -76 | -6 | 5.7 | 25 |

The table shows peak decoding in the searchlight analysis for aLaF (FWE corrected at $p < 0.0000001$ , with an extent threshold of 20 voxels). Results for a more lenient threshold (FWE corrected at $p < 0.05$ , with an extent threshold of 20 voxels) are shown for aLuF. There were no voxels showing significant decoding of the two other attention conditions comprising unattended information (uLaF and uLuF) at this more lenient threshold.

At this high threshold, there was no significant decodable stimulus information in any region when it was not relevant for the task (aLuF, uLaF and uLuf conditions). At a more lenient threshold of p<0.05 FWE corrected, we found coding of unattended feature information of the target object (aLuF) in two small clusters within right temporal lobe and left lateral occipital complex (Table 3). However, there were no significant clusters for relevant or irrelevant features of the distractor (uLaF and uLuF), even at this more lenient threshold.

## 4. Discussion

In this study we used fMRI to understand how spatial and feature attention interact to affect task-related representations in MD regions and the visual cortex. We found that spatial and feature attention interacted multiplicatively. The MD system was highly specific in its representation of task-related information, representing only information about the attended feature of the attended object. We found similar specificity in task-related coding in visual regions. We found no evidence of irrelevant information decoding in MD or visual ROIs, even when that irrelevant information pertained to the target object or task-relevant features.

Our data align with previous inverted encoding model work in fMRI showing stronger and more reliable reconstruction of the task-relevant feature of the target in visual working memory, relative to task-irrelevant feature information or task-relevant features of the distractor (Yu & Shim, 2017). In that work, only the task-relevant feature of the target could be reconstructed in frontal, parietal and occipital cortices during a task where participants were asked to recall either the colour or orientation of one of two presented gratings after a delay (Yu & Shim, 2017). Similarly, in a task where participants reported the colour or orientation of one of three gratings, only the task-relevant feature of the target could be reliably reconstructed from visual areas and IPS (Chen et al., 2021). Interestingly, in this preprint, Chen et al. (2021) found some representation of the irrelevant feature of the target in intraparietal regions, perhaps reflecting attention spreading to both features of the attended grating, but reconstruction of the irrelevant feature was weak and inconsistent. This study only examined task-related representations in occipital and posterior parietal regions. Our study corroborates and extends these studies by demonstrating the dominance of decoding of task-relevant features in frontal and parietal regions, in an attention task with novel objects.

Our findings also extend recent MEG work which reported the multiplicative effect of spatial and feature attention on decoding of object information in this and a similar task (Barnes et al., 2022; Goddard et al., 2022). As well as replicating this effect in a new modality, our results provide spatial information. Our data suggest strong dominance of task-relevant information in MD regions, as predicted for regions strongly engaged in difficult tasks (Assem, Glasser, et al., 2020; Fedorenko et al., 2013) and which are proposed to adjust their responses to carry the most relevant information (Erez & Duncan, 2015; Jackson et al., 2017; Woolgar, Williams, et al., 2015). Moreover, in our searchlight analysis, which is free from *a priori* spatial hypotheses, task-relevant information was most strongly coded in the MD system. However, our data also suggest that the multiplicative effect of these two types of attention is not limited to the MD cortex, but can been seen in widespread areas of the brain including in visual regions, perhaps reflecting the dominance of attended information across the system.

Time-resolved data (e.g., from E/MEG) show initial decoding of all stimulus information (relevant and irrelevant) that is not reflected in our results. In time-resolved data, stimulus information that is ignored (Goddard et al., 2022; Moerel et al., 2021, 2022) or not used to drive behaviour (Robinson et al., 2022) is typically encoded briefly. For instance, Goddard et al. (2022) showed brief initial coding of all stimulus information in occipital regions before resolving to a sustained preferential coding of only task-relevant information from around 400ms. Similarly, other work exploring the temporal dynamics of attention suggest an earlier dominance of task-related representations in frontoparietal regions, that later comes to dominate processing in visual regions (Hebart et al., 2018). The coarse temporal resolution of fMRI may have precluded us from seeing these early neural effects. Thus, we suspect that the widespread and exclusive representation of task-relevant information seen in this study reflects more sustained attentional processes whereby MD regions, and perhaps the neural system more broadly, ultimately prioritise a highly specific task-relevant representation.

These results are seemingly at odds with object-based and feature-based attention theories (e.g., Duncan, 1984; Sàenz et al., 2002). For the former, an object-grouping effect would be expected, such that attention paid to a particular feature of an object facilitates processing of other, unattended, features of the same object. Commensurate with this, behavioural studies indicate better performance when reporting multiple features of a single object than of different objects, even when accounting for spatial selection (Duncan, 1984), and greater interference from distractors in the same object grouping than across different object groupings (e.g., Egly et al., 1994). Neuroimaging work also points to enhanced representation of all attributes of the attended object (Barnes et al., 2022; Jiang et al., 2016; O’Craven et al., 1999; Schoenfeld et al., 2014). In our case, we would expect an object-based effect to be reflected in greater representation of unattended features of the target object, relative to attended and unattended features of the distractor object. Our findings are inconsistent with this prediction: in the MD system, we found no evidence of coding of the unattended feature of the target. However, the whole brain searchlight did reveal coding of the unattended feature of the target in two small clusters of the posterior temporal and lateral occipital regions. Thus, the results in these intermediate regions, typically associated with object processing, are in line with object-based accounts of attention, but, this coding of unattended features did not appear to propagate to high cortical regions. Our data do not support feature-based accounts, whereby a spatially global representation of all attended feature information would be predicted (Jehee et al., 2011), as we found no evidence of neural coding of the attended feature of the distractor.

Recent work by Barnes et al. (2022) provides a lens through which to reconcile the apparent conflict between our results and the predictions stemming from seminal attentional theories. Barnes et al. (2022) utilised a selective attention task that varied in its demands, yielding a more complex picture of the specificity of task-related representations in the brain. The less demanding version of their task resulted in data that fit with predictions of object-based attention, with strong coding of both relevant and irrelevant feature information of the attended object. When discriminations were made more challenging with multiple competing objects, however, the data followed the pattern reported here, with a multiplicative effect of the two attention types. Relatedly, other studies point to weaker representation of distractor information (Zhang & Luck, 2009) and selective prioritisation of attended visual stimuli (Woolgar, Williams, et al., 2015) particularly when stimuli are more difficult to perceive. In line with such observations, load theory (Lavie, 2005; Murphy et al., 2016) posits that load (e.g., increasing perceptual difficulty) affects the degree to which distractors are processed. Together, these findings suggest a potential explanation for why our findings might diverge from what is predicted by object-based accounts: the brain may have different ways of operating depending on the demands of the presenting challenge. Our attention task closely resembled the more demanding attentional task used by Barnes et al. (2022) and, similar to Barnes et al. (2022), we did not see an advantage for the irrelevant feature of the attended object. Our findings may therefore reflect the context of competition between stimulus information in our task, culminating in only task-relevant information being selected for processing.

These considerations mean that a different result may have been obtained if the task were easier. Another consideration is that our decoding analysis involved training and testing on both attend left and right runs, such that the classifier was required to generalise over attended location. While this does not affect the interpretation of our primary result – that representations in MD cortex are highly specific to the task at hand – it would have reduced our sensitivity to detect stimulus-driven representations that are lateralised or retinotopic. Finally, we also cannot rule out that the brain may store irrelevant information differently to relevant information, potentially involving mechanisms that are less visible to neuroimaging (Barbosa et al., 2021; Stokes, 2015).

As a secondary research question, we asked how the representation of task-relevant information was structured in MD regions. We have previously found that coding of attended information in MD regions is stronger when stimuli are physically more similar, and therefore more difficult to perceive (Woolgar et al., 2011; Woolgar, Williams, et al., 2015) (although note (Wen et al., 2018)). This might predict that stimuli closer to the participant’s decision boundary would be the most decodable, in a reversal of the typical distance-to-bound effects observed in visual cortex (Ritchie & Carlson, 2016). Indeed, in our data, pairs of stimuli close to the decision boundary could be discriminated more readily than pairs of stimuli far from the decision boundary, even though the physical difference between stimuli far from the decision boundary was greater. However, rather than being intermediate, medium physical discriminations (i.e., comprising stimuli two feature steps apart) were the most discriminable of all combinations. Therefore, a single axis reflecting the physical discriminability of the stimuli (or its inverse) could not account for the results. Instead, we found that the representation of task-relevant information was captured by two organising axes. One reflected the stimulus categories (e.g., red vs green) and the other, the difficulty of the perceptual decision (easy vs hard). We obtained this result despite modelling out differences in RT at the single trial level (Grinband et al., 2008; Henson, 2007; Todd et al., 2013; Woolgar et al., 2014). Our findings here fit within a body of literature showing that prefrontal cortex (PFC) responses reflect the behavioural relevance of stimuli as much or more than their physical properties (e.g., Cromer et al., 2010; Erez & Duncan, 2015; Kadohisa et al., 2013; Sakagami & Niki, 1994; Wisniewski et al., 2023), and work emphasising high-dimensionality in PFC patterns that are related to behaviour (Badre et al., 2021; Bartolo et al., 2020; Rigotti et al., 2013). Our data add to this picture by confirming that MD responses reflect both an abstracted representation of difficulty that generalised over stimulus information, and representation of categorical colour information (which in this case equates with decision, although not motor response), generalised over differences in difficulty.

### 4.1. Conclusions

In this study we used fMRI to understand how spatial and feature attention interact to affect task-related representations in MD regions and elsewhere in the brain. Our data strongly suggest that in the context of interacting spatial and feature attention in a difficult perceptual categorisation task, the MD system is highly specific in its representation of task-related information, representing only information at the intersection of the two types of attention. We also found that MD representations of the selected information reflected both categorical task-relevant stimulus decisions and task difficulty. Finally, we found that while the effect was strongest in MD regions, the multiplicative attention effect was widespread throughout the brain, and applied even to early visual regions.

In summary, our data shed light on the interacting effects of spatial and feature attention, a context more closely resembling the way different subtypes of attention are used in real life. Rather than boosting processing of whole objects or relevant features across space, as might be predicted when examining the effects of spatial and feature attention alone, neural activity in our data appeared to reflect all-or-nothing tuning to behaviourally relevant information. These results emphasise the importance of considering interacting attentional demands to understand the mechanisms underpinning selective attention.

## 5. Supplementary Materials

### 5.1. Behavioural accuracy and reaction time

**Figure S1.**
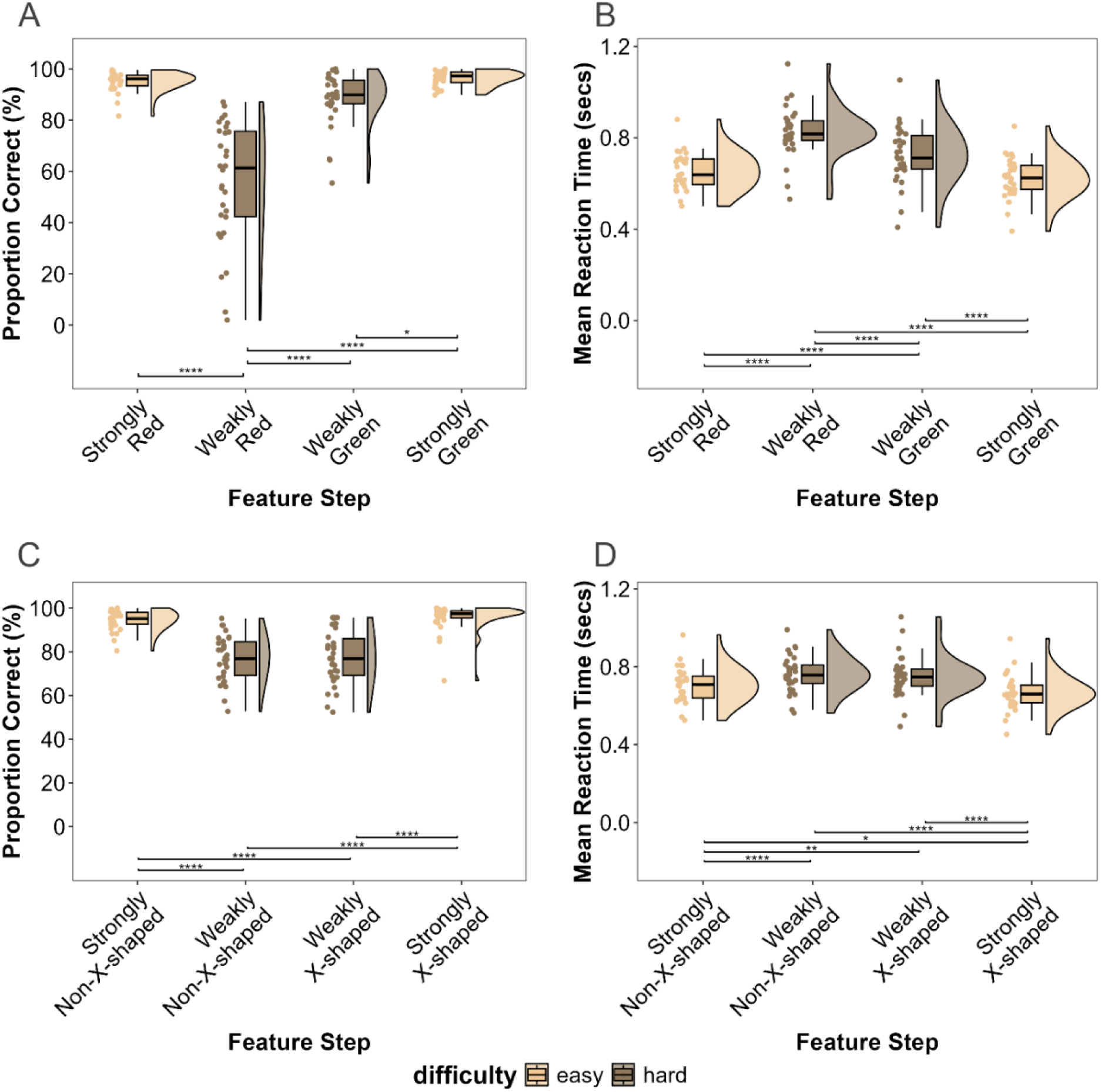
Panels A and B: Proportion correct by relevant feature value, for colour (A) and shape (B) trials. Panels C and D: Average reaction time (in seconds) for correct trials across all participants, across relevant feature steps, for colour (C) and shape (D) trials. Results are coloured according to whether stimuli were ‘easy’ (i.e., far from the decision boundary) or ‘hard’ (i.e., close to the decision boundary). Scatter at left depicts mean accuracy or mean reaction time for the given task for each participant. Boxplots indicate mean accuracy or mean reaction time (central line), box hinges indicate first and third quartiles, and box whiskers range from hinge to largest and smallest decoding accuracy value within 1.5 x applicable quartile range. Data distribution is indicated at right for each condition (half violin). Significant pairwise comparisons marked with Bonferroni adjusted p-value codes as follows: * < 0.05, ** < 0.01, *** < 0.001, **** < 0.0001.

### 5.2. MD decoding of attended and unattended stimulus information, shown for colour and shape information separately

**Figure S2.**
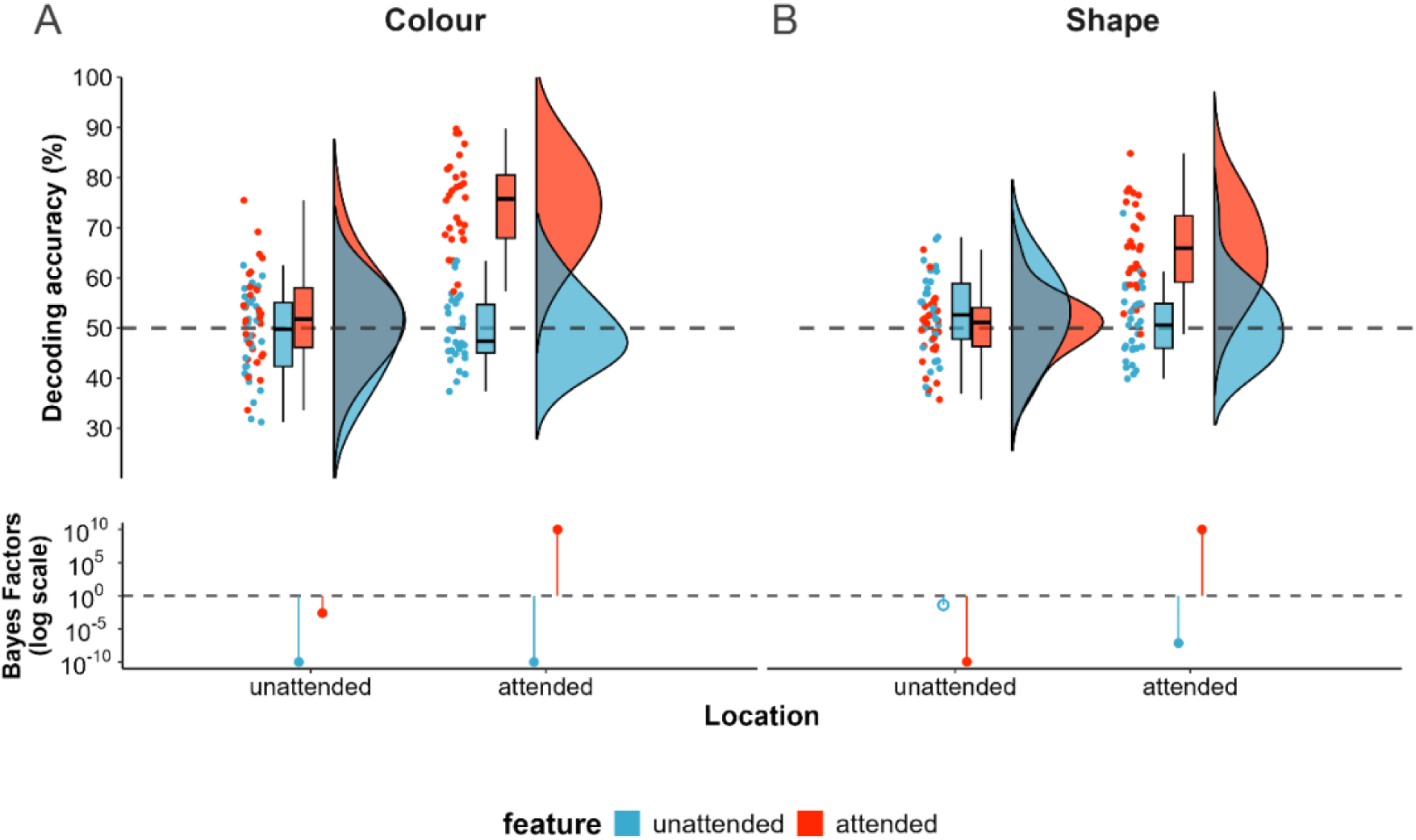
The effect of two interacting subtypes of attention on decoding of feature information for averaged MD regions, shown separately for colour (Panel A) or shape (Panel B) information. Decoding accuracy reflects average accuracy in decoding colour or shape information in the object that was in the unattended (left plots) or attended (right plots) location, when the feature in question was attended (red) or unattended (blue). Dots depict individual subject results, boxplots indicate median (central line), first and third quartiles (box hinges), and largest and smallest decoding accuracy value within 1.5 x applicable quartile range (whiskers), half violin depicts data distribution. The strength of evidence for above-chance (H1) or at-chance (H0) decoding was quantified using Bayesian t-tests and is shown as Bayes Factors for each of the conditions on a logarithmic scale, with solid circles reflecting at least moderate evidence in favour of H1 or H0, and open circles reflecting evidence was anecdotal or inconclusive.

### 5.3. MD decoding by stimulus discriminability, after restricting analysis to those contrast that straddle the participant’s decision boundary

**Figure S3.**
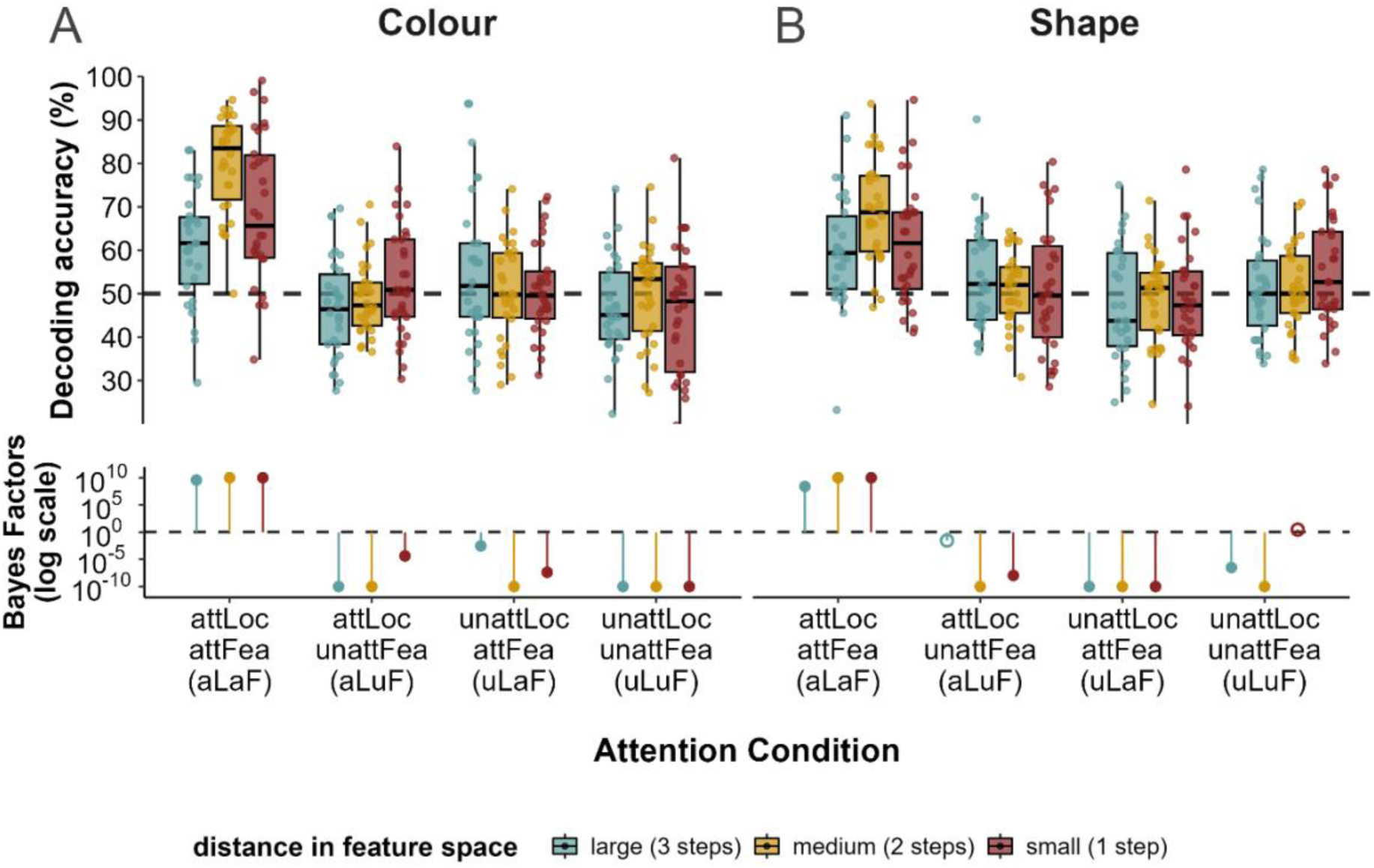
Decoding of colour (Panel A) and shape (Panel B) information in each of the spatial and feature attention condition for each distance in feature space level, collapsed across all MD regions. In contrast to Figure 3, distance in feature space comparisons in this plot comprise only comparisons that cross the decision boundary. This represents a difference for the small distances in feature space level (maroon bars) only. That is, in this plot, large distances in feature space comprise three feature steps (pairwise comparison 1v4), medium distances in feature space comprise two feature steps (pairwise comparisons 1v3, 2v4), and small distances in feature space comprise one feature step (pairwise comparison 2v3). Dots depict individual subject results, boxplots indicate median (central line), first and third quartiles (box hinges), and largest and smallest decoding accuracy value within 1.5 x applicable quartile range (whiskers). The strength of evidence for above-chance (H1) or at-chance (H0) decoding was quantified using Bayesian t-tests and is shown as Bayes Factors for each of the conditions on a logarithmic scale, with solid circles reflecting at least moderate evidence in favour of H1 or H0, and open circles reflecting evidence was anecdotal or inconclusive.

### 5.4. Visual cortex decoding by stimulus discriminability, after restricting analysis to those contrast that straddle the participant’s decision boundary

**Figure S4.**
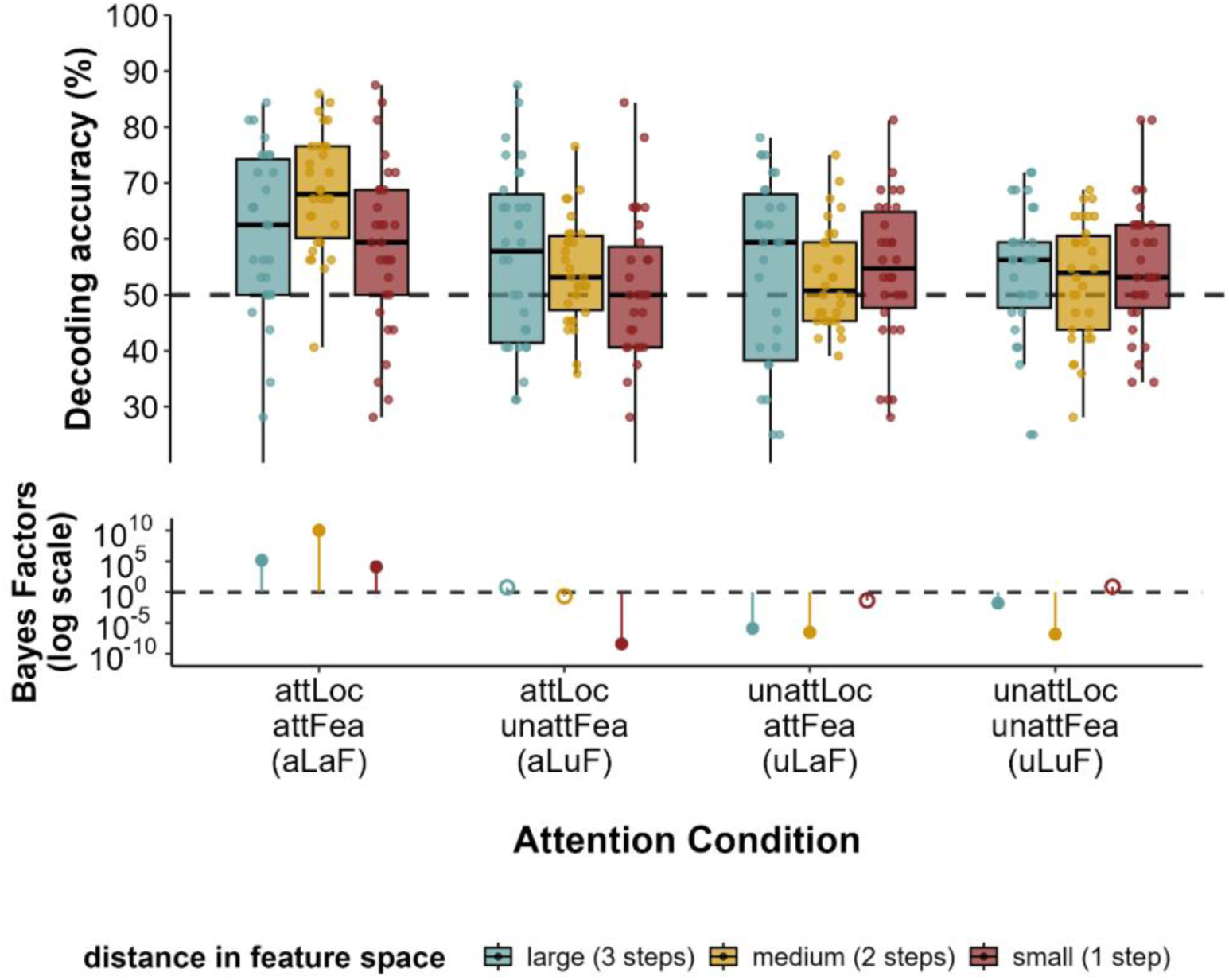
Decoding of stimulus information (averaged over colour and shape information, as not significantly different between these types of information) for visual cortex (BA17/18). In contrast to Figure 6, distance in feature space comparisons in this plot comprise only comparisons that cross the decision boundary. This represents a difference for the small distances in feature space level (maroon bars) only. That is, in this plot, large distances in feature space comprise three feature steps (pairwise comparison 1v4), medium distances in feature space comprise two feature steps (pairwise comparisons 1v3, 2v4), and small distances in feature space comprise one feature step (pairwise comparison 2v3). Dots depict individual subject results, boxplots indicate median (central line), first and third quartiles (box hinges), and largest and smallest decoding accuracy value within 1.5 x applicable quartile range (whiskers). The strength of evidence for above-chance (H1) or at-chance (H0) decoding was quantified using Bayesian t-tests and is shown as Bayes Factors for each of the conditions on a logarithmic scale, with solid circles reflecting at least moderate evidence in favour of H1 or H0, and open circles reflecting evidence was anecdotal or inconclusive.

## 6. CRediT authorship contribution statement

**Nadene Dermody**: Conceptualization, Data curation, Formal analysis, Writing – original draft, Writing – review and editing

**Romy Lorenz**: Conceptualization, Formal analysis, Supervision, Writing – review and editing

**Erin Goddard**: Conceptualization, Formal analysis, Writing – review and editing

**Arno Villringer**: Funding acquisition, Resources, Supervision

**Alexandra Woolgar**: Conceptualization, Formal analysis, Funding acquisition, Supervision, Writing – review and editing

## Acknowledgements

We thank Alice Hodapp and Paul Steinfath for their help in collecting data for this study, and Jöran Lepsein for technical support.

## References

Arazi, A., Yeshurun, Y., & Dinstein, I. (2019). Neural Variability Is Quenched by Attention. Journal of Neuroscience, 39(30), 5975–5985. 10.1523/JNEUROSCI.0355-19.2019

Assem, M., Blank, I. A., Mineroff, Z., Ademoğlu, A., & Fedorenko, E. (2020). Activity in the fronto-parietal multiple-demand network is robustly associated with individual differences in working memory and fluid intelligence. Cortex, 131, 1–16. 10.1016/J.CORTEX.2020.06.013

Assem, M., Glasser, M. F., Van Essen, D. C., & Duncan, J. (2020). A Domain-General Cognitive Core Defined in Multimodally Parcellated Human Cortex. Cerebral Cortex, 30(8), 4361– 4380. 10.1093/cercor/bhaa023

Badre, D., Bhandari, A., Keglovits, H., & Kikumoto, A. (2021). The dimensionality of neural representations for control. Current Opinion in Behavioral Sciences, 38, 20–28. 10.1016/j.cobeha.2020.07.002

Barbosa, J., Lozano-Soldevilla, D., & Compte, A. (2021). Pinging the brain with visual impulses reveals electrically active, not activity-silent, working memories. PLOS Biology, 19(10), e3001436. 10.1371/journal.pbio.3001436

Barnes, L., Goddard, E., & Woolgar, A. (2022). Neural Coding of Visual Objects Rapidly Reconfigures to Reflect Subtrial Shifts in Attentional Focus. Journal of Cognitive Neuroscience, 34(5), 806–822. 10.1162/jocn_a_01832

Bartolo, R., Saunders, R. C., Mitz, A. R., & Averbeck, B. B. (2020). Dimensionality, information and learning in prefrontal cortex. PLOS Computational Biology, 16(4), e1007514. 10.1371/journal.pcbi.1007514

Bates, D., Mächler, M., Bolker, B., & Walker, S. (2015). Fitting Linear Mixed-Effects Models Using lme4. Journal of Statistical Software, 67, 1–48. 10.18637/jss.v067.i01

Battistoni, E., Kaiser, D., Hickey, C., & Peelen, M. V. (2020). The time course of spatial attention during naturalistic visual search. Cortex, 122, 225–234. 10.1016/j.cortex.2018.11.018

Brawn, P. T., & Snowden, R. J. (2000). Attention to overlapping objects: Detection and discrimination of luminance changes. Journal of Experimental Psychology: Human Perception and Performance, 26(1), 342–358. 10.1037/0096-1523.26.1.342

Burrows, B. E., & Moore, T. (2009). Influence and Limitations of Popout in the Selection of Salient Visual Stimuli by Area V4 Neurons. Journal of Neuroscience, 29(48), 15169–15177. 10.1523/JNEUROSCI.3710-09.2009

Carrasco, M. (2011). Visual attention: The past 25 years. Vision Research, 51(13), 1484–1525. 10.1016/J.VISRES.2011.04.012

Chang, C.-C., & Lin, C.-J. (2011). LIBSVM: A library for support vector machines. ACM Transactions on Intelligent Systems and Technology, 2(3), 27:1-27:27. 10.1145/1961189.1961199

Chen, J., Scotti, P. S., Dowd, E. W., & Golomb, J. D. (2021). Neural Representations of Task-relevant and Task-irrelevant Features of Attended Objects. bioRxiv, 2021.05.21.445168. 10.1101/2021.05.21.445168

Chen, X., Hoffmann, K.-P., Albright, T. D., & Thiele, A. (2012). Effect of feature-selective attention on neuronal responses in macaque area MT. Journal of Neurophysiology, 107(5), 1530–1543.

Cohen, M. R., & Maunsell, J. H. R. (2009). Attention improves performance primarily by reducing interneuronal correlations. Nature Neuroscience, 12(12), 1594–1600. 10.1038/nn.2439

Cole, M. W., Reynolds, J. R., Power, J. D., Repovs, G., Anticevic, A., & Braver, T. S. (2013). Multi-task connectivity reveals flexible hubs for adaptive task control. Nature Neuroscience, 16(9), 1348–1355. 10.1038/nn.3470

Cole, M. W., & Schneider, W. (2007). The cognitive control network: Integrated cortical regions with dissociable functions. NeuroImage, 37(1), 343–360. 10.1016/J.NEUROIMAGE.2007.03.071

Cromer, J. A., Roy, J. E., & Miller, E. K. (2010). Representation of multiple, independent categories in the primate prefrontal cortex. Neuron, 66(5), 796–807.

Desimone, R., & Duncan, J. (1995). Neural mechanisms of selective visual attention. Annual Review of Neuroscience, 18, 193–222. 10.1146/annurev.ne.18.030195.001205

Duncan, J. (1984). Selective attention and the organization of visual information. Journal of Experimental Psychology. General, 113(4), 501–517.

Duncan, J. (2001). An adaptive coding model of neural function in prefrontal cortex. Nature Reviews. Neuroscience, 2(11), 820–829. 10.1038/35097575

Duncan, J. (2010). The multiple-demand (MD) system of the primate brain: Mental programs for intelligent behaviour. Trends in Cognitive Sciences, 14(4), 172–179. 10.1016/j.tics.2010.01.004

Duncan, J., & Owen, A. M. (2000). Common regions of the human frontal lobe recruited by diverse cognitive demands. Trends in Neurosciences, 23(10), 475–483. 10.1016/S0166-2236(00)01633-7

Egly, R., Driver, J., & Rafal, R. D. (1994). Shifting Visual Attention Between Objects and Locations: Evidence From Normal and Parietal Lesion Subjects. Journal of Experimental Psychology: General, 123(2), 161–177. 10.1037/0096-3445.123.2.161

Erez, Y., & Duncan, J. (2015). Discrimination of Visual Categories Based on Behavioral Relevance in Widespread Regions of Frontoparietal Cortex. Journal of Neuroscience, 35(36), 12383– 12393. 10.1523/JNEUROSCI.1134-15.2015

Fedorenko, E., Duncan, J., & Kanwisher, N. (2013). Broad domain generality in focal regions of frontal and parietal cortex. Proceedings of the National Academy of Sciences of the United States of America, 110(41), 16616–16621. 10.1073/pnas.1315235110

Fox, M. D., Snyder, A. Z., Vincent, J. L., Corbetta, M., Van Essen, D. C., & Raichle, M. E. (2005). The human brain is intrinsically organized into dynamic, anticorrelated functional networks. Proceedings of the National Academy of Sciences, 102(27), 9673–9678. 10.1073/pnas.0504136102

Goddard, E., Carlson, T. A., & Woolgar, A. (2022). Spatial and Feature-selective Attention Have Distinct, Interacting Effects on Population-level Tuning. Journal of Cognitive Neuroscience, 34(2), 290–312. 10.1162/JOCN_A_01796

Gratton, C., Sun, H., & Petersen, S. E. (2018). Control networks and hubs. Psychophysiology, 55(3), e13032. 10.1111/psyp.13032

Grinband, J., Wager, T. D., Lindquist, M., Ferrera, V. P., & Hirsch, J. (2008). Detection of time-varying signals in event-related fMRI designs. NeuroImage, 43(3), 509–520. 10.1016/j.neuroimage.2008.07.065

Grootswagers, T., Robinson, A. K., Shatek, S. M., & Carlson, T. A. (2021). The neural dynamics underlying prioritisation of task-relevant information. *Neurons, Behavior*, Data Analysis, and Theory, 5(1), 1–17. 10.51628/001c.21174

Hebart, M. N., Bankson, B. B., Harel, A., Baker, C. I., & Cichy, R. M. (2018). The representational dynamics of task and object processing in humans. eLife, 7, e32816. 10.7554/eLife.32816

Hebart, M. N., Görgen, K., & Haynes, J.-D. (2015). The Decoding Toolbox (TDT): A versatile software package for multivariate analyses of functional imaging data. Frontiers in Neuroinformatics, 8. https://www.frontiersin.org/articles/10.3389/fninf.2014.00088

Heinze, H. J., Mangun, G. R., Burchert, W., Hinrichs, H., Scholz, M., Münte, T. F., Gös, A., Scherg, M., Johannes, S., Hundeshagen, H., Gazzaniga, M. S., & Hillyard, S. A. (1994). Combined spatial and temporal imaging of brain activity during visual selective attention in humans. Nature, 372(6506), 543–546. 10.1038/372543a0

Henson, R. (2007). Efficient Experimental Design for fMRI. In Statistical Parametric Mapping (pp. 193–210). Elsevier. 10.1016/B978-012372560-8/50015-2

Jackson, J. B., Feredoes, E., Rich, A. N., Lindner, M., & Woolgar, A. (2021). Concurrent neuroimaging and neurostimulation reveals a causal role for dlPFC in coding of task-relevant information. Communications Biology, 4(1), 1–16. 10.1038/s42003-021-02109-x

Jackson, J. B., Rich, A. N., Williams, M. A., & Woolgar, A. (2017). Feature-selective Attention in Frontoparietal Cortex: Multivoxel Codes Adjust to Prioritize Task-relevant Information. Journal of Cognitive Neuroscience, 29(2), 310–321. 10.1162/jocn_a_01039

Jackson, J. B., & Woolgar, A. (2018). Adaptive coding in the human brain: Distinct object features are encoded by overlapping voxels in frontoparietal cortex. Cortex, 108, 25–34. 10.1016/j.cortex.2018.07.006

Jeffreys, H. (1998). The theory of probability. OuP Oxford.

Jehee, J. F. M., Brady, D. K., & Tong, F. (2011). Attention Improves Encoding of Task-Relevant Features in the Human Visual Cortex. Journal of Neuroscience, 31(22), 8210–8219. 10.1523/JNEUROSCI.6153-09.2011

Jiang, J., Summerfield, C., & Egner, T. (2016). Visual Prediction Error Spreads Across Object Features in Human Visual Cortex. Journal of Neuroscience, 36(50), 12746–12763. 10.1523/JNEUROSCI.1546-16.2016

Kadohisa, M., Petrov, P., Stokes, M., Sigala, N., Buckley, M., Gaffan, D., Kusunoki, M., & Duncan, J. (2013). Dynamic construction of a coherent attentional state in a prefrontal cell population. Neuron, 80(1), 235–246. 10.1016/j.neuron.2013.07.041

Kanashiro, T., Ocker, G. K., Cohen, M. R., & Doiron, B. (2017). Attentional modulation of neuronal variability in circuit models of cortex. eLife, 6, e23978. 10.7554/eLife.23978

Kastner, S., Pinsk, M. A., De Weerd, P., Desimone, R., & Ungerleider, L. G. (1999). Increased Activity in Human Visual Cortex during Directed Attention in the Absence of Visual Stimulation. Neuron, 22(4), 751–761. 10.1016/S0896-6273(00)80734-5

Keller, A. S., Jagadeesh, A. V., Bugatus, L., Williams, L. M., & Grill-Spector, K. (2022). Attention enhances category representations across the brain with strengthened residual correlations to ventral temporal cortex. NeuroImage, 249, 118900. 10.1016/j.neuroimage.2022.118900

Kriegeskorte, N., Goebel, R., & Bandettini, P. (2006). Information-based functional brain mapping. Proceedings of the National Academy of Sciences, 103(10), 3863–3868. 10.1073/pnas.0600244103

Kuznetsova, A., Brockhoff, P. B., & Christensen, R. H. B. (2017). lmerTest Package: Tests in Linear Mixed Effects Models. Journal of Statistical Software, 82(13), 1–26. 10.18637/jss.v082.i13

Lavie, N. (2005). Distracted and confused?: Selective attention under load. Trends in Cognitive Sciences, 9(2), 75–82. 10.1016/j.tics.2004.12.004

Li, Y., Wang, F., Chen, Y., Cichocki, A., & Sejnowski, T. (2018). The Effects of Audiovisual Inputs on Solving the Cocktail Party Problem in the Human Brain: An fMRI Study. Cerebral Cortex, 28(10), 3623–3637. 10.1093/cercor/bhx235

Liu, T. (2019). Feature-based attention: Effects and control. Current Opinion in Psychology, 29, 187–192. 10.1016/j.copsyc.2019.03.013

Liu, T., Stevens, S. T., & Carrasco, M. (2007). Comparing the time course and efficacy of spatial and feature-based attention. Vision Research, 47(1), 108–113. 10.1016/j.visres.2006.09.017

Lu, R., Michael, E., Scrivener, C. L., Jackson, J. B., Duncan, J., & Woolgar, A. (2024). Parietal alpha stimulation causally enhances attentional information coding in evoked and oscillatory activity (p. 2023.11.14.567111). bioRxiv. 10.1101/2023.11.14.567111

Luck, S. J., Chelazzi, L., Hillyard, S. A., & Desimone, R. (1997). Neural mechanisms of spatial selective attention in areas V1, V2, and V4 of macaque visual cortex. Journal of Neurophysiology, 77(1), 24–42. 10.1152/JN.1997.77.1.24/ASSET/IMAGES/LARGE/JNP.JA28F16.JPEG

Luo, T. Z., & Maunsell, J. H. R. (2018). Attentional Changes in Either Criterion or Sensitivity Are Associated with Robust Modulations in Lateral Prefrontal Cortex. Neuron, 97(6), 1382–1393.e7. 10.1016/j.neuron.2018.02.007

Maunsell, J. H. R., & Treue, S. (2006). Feature-based attention in visual cortex. Trends in Neurosciences, 29(6), 317–322. 10.1016/j.tins.2006.04.001

McAdams, C. J., & Maunsell, J. H. R. (1999). Effects of Attention on Orientation-Tuning Functions of Single Neurons in Macaque Cortical Area V4. Journal of Neuroscience, 19(1), 431–441. 10.1523/JNEUROSCI.19-01-00431.1999

Meteyard, L., & Davies, R. A. I. (2020). Best practice guidance for linear mixed-effects models in psychological science. Journal of Memory and Language, 112, 104092. 10.1016/j.jml.2020.104092

Miller, E. K., & Cohen, J. D. (2001). An integrative theory of prefrontal cortex function. Annual Review of Neuroscience, 24, 167–202. 10.1146/annurev.neuro.24.1.167

Mitchell, J. F., Sundberg, K. A., & Reynolds, J. H. (2007). Differential Attention-Dependent Response Modulation across Cell Classes in Macaque Visual Area V4. Neuron, 55(1), 131–141. 10.1016/j.neuron.2007.06.018

Moerel, D., Grootswagers, T., Robinson, A. K., Shatek, S. M., Woolgar, A., Carlson, T. A., & Rich, A. N. (2022). The time-course of feature-based attention effects dissociated from temporal expectation and target-related processes. Scientific Reports, 12(1), 6968. 10.1038/s41598-022-10687-x

Moerel, D., Rich, A. N., & Woolgar, A. (2021). Selective attention and decision-making have separable neural bases in space and time [Preprint]. Neuroscience. 10.1101/2021.02.28.433294

Moran, J., & Desimone, R. (1985). Selective Attention Gates Visual Processing in the Extrastriate Cortex. Science, 229(4715), 782–784. 10.1126/SCIENCE.4023713

Morey, R., & Rouder, J. (2011). Bayes Factor Approaches for Testing Interval Null Hypotheses. Psychological Methods, 16, 406–419. 10.1037/a0024377

Morey, R., & Rouder, J. (2018). BayesFactor: Computation of Bayes Factors for Common Designs. https://CRAN.R-project.org/package=BayesFactor

Murphy, G., Groeger, J. A., & Greene, C. M. (2016). Twenty years of load theory—Where are we now, and where should we go next? Psychonomic Bulletin & Review, 23(5), 1316–1340. 10.3758/s13423-015-0982-5

Noudoost, B., Chang, M. H., Steinmetz, N. A., & Moore, T. (2010). Top-down control of visual attention. Current Opinion in Neurobiology, 20(2), 183–190. 10.1016/j.conb.2010.02.003

O’Connor, D. H., Fukui, M. M., Pinsk, M. A., & Kastner, S. (2002). Attention modulates responses in the human lateral geniculate nucleus. Nature Neuroscience, 5(11), 1203– 1209. 10.1038/nn957

O’Craven, K. M., Downing, P. E., & Kanwisher, N. (1999). fMRI evidence for objects as the units of attentional selection. Nature 1999 *401*:6753, *401*(6753), 584–587. 10.1038/44134

Olszowy, W., Aston, J., Rua, C., & Williams, G. B. (2019). Accurate autocorrelation modeling substantially improves fMRI reliability. Nature Communications, 10(1), 1220. 10.1038/s41467-019-09230-w

Reynolds, J. H., & Heeger, D. J. (2009). The normalization model of attention. Neuron, 61(2), 168–185. 10.1016/j.neuron.2009.01.002

Rigotti, M., Barak, O., Warden, M. R., Wang, X.-J., Daw, N. D., Miller, E. K., & Fusi, S. (2013). The importance of mixed selectivity in complex cognitive tasks. Nature, 497(7451), 585– 590.

Ritchie, J. B., & Carlson, T. A. (2016). Neural Decoding and “Inner” Psychophysics: A Distance-to-Bound Approach for Linking Mind, Brain, and Behavior. Frontiers in Neuroscience, 10. https://www.frontiersin.org/articles/10.3389/fnins.2016.00190

Robinson, A. K., Rich, A. N., & Woolgar, A. (2022). Linking the Brain with Behavior: The Neural Dynamics of Success and Failure in Goal-directed Behavior. Journal of Cognitive Neuroscience, 34(4), 639–654. 10.1162/jocn_a_01818

Rossi, A. F., & Paradiso, M. A. (1995). Feature-specific effects of selective visual attention. Vision Research, 35(5), 621–634. 10.1016/0042-6989(94)00156-G

Rouder, J. N., Speckman, P. L., Sun, D., Morey, R. D., & Iverson, G. (2009). Bayesian t tests for accepting and rejecting the null hypothesis. Psychonomic Bulletin & Review, 16(2), 225–237. 10.3758/PBR.16.2.225

Sàenz, M., Buraĉas, G. T., & Boynton, G. M. (2002). Global effects of feature-based attention in human visual cortex. Nature Neuroscience 2002 5:7, 5(7), 631–632. 10.1038/nn876

Sàenz, M., Buraĉas, G. T., & Boynton, G. M. (2003). Global feature-based attention for motion and color. Vision Research, 43(6), 629–637. 10.1016/S0042-6989(02)00595-3

Sakagami, M., & Niki, H. (1994). Encoding of behavioral significance of visual stimuli by primate prefrontal neurons: Relation to relevant task conditions. Experimental Brain Research, 97(3), 423–436. 10.1007/BF00241536

Schoenfeld, M. A., Hopf, J.-M., Merkel, C., Heinze, H.-J., & Hillyard, S. A. (2014). Object-based attention involves the sequential activation of feature-specific cortical modules. Nature Neuroscience, 17(4), Article 4. 10.1038/nn.3656

Schultz, D. H., Ito, T., & Cole, M. W. (2022). Global connectivity fingerprints predict the domain generality of multiple-demand regions. Cerebral Cortex, 32(20), 4464–4479. 10.1093/cercor/bhab495

Seeley, W. W., Menon, V., Schatzberg, A. F., Keller, J., Glover, G. H., Kenna, H., Reiss, A. L., & Greicius, M. D. (2007). Dissociable Intrinsic Connectivity Networks for Salience Processing and Executive Control. Journal of Neuroscience, 27(9), 2349–2356. 10.1523/JNEUROSCI.5587-06.2007

Stokes, M. G. (2015). ‘Activity-silent’ working memory in prefrontal cortex: A dynamic coding framework. Trends in Cognitive Sciences, 19(7), 394–405. 10.1016/j.tics.2015.05.004

Teichmann, L., Moerel, D., Baker, C., & Grootswagers, T. (2021). An empirically-driven guide on using Bayes Factors for M/EEG decoding. bioRxiv, 2021.06.23.449663. 10.1101/2021.06.23.449663

Thiele, A., & Bellgrove, M. A. (2018). Neuromodulation of Attention. Neuron, 97(4), 769–785. 10.1016/j.neuron.2018.01.008

Thiele, A., Brandt, C., Dasilva, M., Gotthardt, S., Chicharro, D., Panzeri, S., & Distler, C. (2016). Attention Induced Gain Stabilization in Broad and Narrow-Spiking Cells in the Frontal Eye-Field of Macaque Monkeys. Journal of Neuroscience, 36(29), 7601–7612. 10.1523/JNEUROSCI.0872-16.2016

Todd, M. T., Nystrom, L. E., & Cohen, J. D. (2013). Confounds in multivariate pattern analysis: Theory and rule representation case study. NeuroImage, 77, 157–165. 10.1016/j.neuroimage.2013.03.039

Treue, S., & Martínez-Trujillo, J. C. (1999). Feature-based attention influences motion processing gain in macaque visual cortex. Nature, 399(6736), 575–579. 10.1038/21176

Vincent, J. L., Kahn, I., Snyder, A. Z., Raichle, M. E., & Buckner, R. L. (2008). Evidence for a Frontoparietal Control System Revealed by Intrinsic Functional Connectivity. Journal of Neurophysiology, 100(6), 3328–3342. 10.1152/jn.90355.2008

Wagenmakers, E.-J., Love, J., Marsman, M., Jamil, T., Ly, A., Verhagen, J., Selker, R., Gronau, Q. F., Dropmann, D., Boutin, B., Meerhoff, F., Knight, P., Raj, A., van Kesteren, E.-J., van Doorn, J., Šmíra, M., Epskamp, S., Etz, A., Matzke, D., … Morey, R. D. (2018). Bayesian inference for psychology. Part II: Example applications with JASP. Psychonomic Bulletin & Review, 25(1), 58–76. 10.3758/s13423-017-1323-7

Wen, T., Mitchell, D. J., & Duncan, J. (2018). Response of the multiple-demand network during simple stimulus discriminations. NeuroImage, 177, 79–87. 10.1016/j.neuroimage.2018.05.019

Wetzels, R., Matzke, D., Lee, M. D., Rouder, J. N., Iverson, G. J., & Wagenmakers, E.-J. (2011). Statistical Evidence in Experimental Psychology: An Empirical Comparison Using 855 t Tests. Perspectives on Psychological Science, 6(3), 291–298. 10.1177/1745691611406923

Wisniewski, D., González-García, C., Formica, S., Woolgar, A., & Brass, M. (2023). Adaptive coding of stimulus information in human frontoparietal cortex during visual classification. NeuroImage, 274, 120150. 10.1016/j.neuroimage.2023.120150

Woolgar, A., Afshar, S., Williams, M. A., & Rich, A. N. (2015). Flexible coding of task rules in frontoparietal cortex: An adaptive system for flexible cognitive control. Journal of Cognitive Neuroscience, 27(10), 1895–1911. 10.1162/jocn_a_00827

Woolgar, A., Golland, P., & Bode, S. (2014). Coping with confounds in multivoxel pattern analysis: What should we do about reaction time differences? A comment on Todd, Nystrom & Cohen 2013. Neuroimage, 98, 506–512. 10.1016/j.neuroimage.2014.04.059

Woolgar, A., Hampshire, A., Thompson, R., & Duncan, J. (2011). Adaptive Coding of Task-Relevant Information in Human Frontoparietal Cortex. Journal of Neuroscience, 31(41), 14592–14599. 10.1523/JNEUROSCI.2616-11.2011

Woolgar, A., Jackson, J. B., & Duncan, J. (2016). Coding of visual, auditory, rule, and response information in the brain: 10 years of multivoxel pattern analysis. Journal of Cognitive Neuroscience, 28(10), 1433–1454. 10.1162/jocn_a_00981

Woolgar, A., Williams, M. A., & Rich, A. N. (2015). Attention enhances multi-voxel representation of novel objects in frontal, parietal and visual cortices. Neuroimage, 109, 429–437. 10.1016/j.neuroimage.2014.12.083

Yu, Q., & Shim, W. M. (2017). Occipital, parietal, and frontal cortices selectively maintain task-relevant features of multi-feature objects in visual working memory. NeuroImage, 157, 97–107. 10.1016/J.NEUROIMAGE.2017.05.055

Zhang, W., & Luck, S. J. (2009). Feature-based attention modulates feedforward visual processing. Nature Neuroscience, 12(1), Article 1. 10.1038/nn.2223

Zheng, Y., Lu, R., & Woolgar, A. (2024). Radical flexibility of neural representation in frontoparietal cortex and the challenge of linking it to behaviour. Current Opinion in Behavioral Sciences, 57, 101392. 10.1016/j.cobeha.2024.101392

